# CDK4/6 inhibition induces a DNA damage-independent senescence-associated secretory phenotype driven by delayed activation of NF-κB

**DOI:** 10.1101/2025.08.25.672139

**Authors:** Joanna Lan-Hing Yeung, Justin Rendleman, Lauren Anderson Westcott, Arnold Ou, Matthew Pressler, Nicole Pagane, Irene Duba, Ria Hosuru, Bat-Ider Tumenbayar, Viviana I. Risca

## Abstract

Cellular senescence consists of regulated cell phenotypes associated with permanent exit from the cell cycle in response to stressors such as genomic instability. The consequences of senescence go beyond individual cells due to the senescence associated secretory phenotype (SASP), which can induce inflammation in neighboring cells. Some cancers respond to CDK4/6 inhibitors (CDK4/6i)—a family of targeted therapies that inhibit proliferation—with a senescence-like phenotype in the absence of DNA damage. We asked how the SASP and the transcriptional regulatory profile triggered by CDK4/6i-driven arrest compares to the canonical NF-κB-regulated SASP triggered by DNA damage. We profiled the temporal dynamics of transcriptional regulation in response to the CDK4/6i, palbociclib, and the DNA damaging agent, doxorubicin. We found that, although upregulation of NF-κB driven-SASP genes is shared across both drugs, it is delayed in CDK4/6i. This coincides with slower enhancer activation and epigenetic changes. Interestingly, ATM/ATR inhibition does not affect CDK4/6i-induced NF-κB nuclear localization, pointing to an alternative mechanism driving NF-κB activity in the absence of DNA damage. Inhibiting NF-κB suppresses the expression of shared SASP genes without reversing stable arrest. This points to SASP manipulation as a potential therapeutic strategy, and resolves an ongoing controversy about the nature of cell cycle arrest-driven SASP.

## INTRODUCTION

Cellular senescence is a state of stable cell cycle arrest that suppresses tumorigenesis by limiting the proliferation of cells that are aged, damaged, or stressed (Ajoolabady et al. 2025). However, even in established tumors, certain therapeutics can force cancer cells into therapy-induced senescence and thereby curb tumor growth (Schmitt et al. 2022). While DNA damage has historically been considered a requirement for entry into senescence, emerging targeted therapies can induce senescent-like states outside of DNA damaging agents and radiation; these include inhibitors of the cyclin-dependent kinases CDK4 and CDK6 (CDK4/6i) (Kovatcheva et al. 2017; Klein et al. 2018; Wagner and Gil 2020). An outstanding question in the field is whether senescence, when caused by dramatically different triggers, converges on a common cell state, or represents a heterogeneous set of states sharing core phenotypic qualities. It is imperative to understand the molecular mechanisms underlying distinct growth-arrest states, as regulatory networks intrinsic to normal aging and stress responses can ultimately shape therapeutic outcomes (Lee and Schmitt 2019; Schmitt et al. 2022; Gleason et al. 2024).

Although senescent cells can no longer divide, they remain metabolically active and acquire features that continually evolve (Schmitt et al. 2022). As such, no universal biomarker of senescence exists, and identification of senescent cells relies on the characterization of several hallmarks (Hernandez-Segura et al. 2018). Beyond stable growth arrest, senescent cells exhibit enlarged morphology, heightened lysosomal content to maintain homeostasis in stressed conditions, increased nuclear size, as well as extensive alterations in the chromatin landscape, including senescence-associated heterochromatic foci (SAHFs) (Hernandez-Segura et al. 2018). Importantly, senescent cells secrete a complex network of cytokines and chemokines, along with other bioactive factors, known as the senescence-associated secretory phenotype (SASP) (Wang et al. 2024); this process increases the paracrine signaling of these “dormant” cells, thus imbuing them with significant influence over the neighboring environment.

Given its effects on the tumor microenvironment, identifying regulators of the SASP across distinct therapeutic triggers of senescence is critical for understanding its impact on disease outcomes. The SASP acts as a double-edged sword, having pro- and anti-tumorigenic effects, depending on its composition (Lau and David 2019). The benefits of the SASP include reinforcing stable arrest through both autocrine and paracrine signals, while facilitating a robust anti-tumor immune response. In contrast, a chronic pro-inflammatory SASP drives increased invasive and migratory capacity of cancer cells, promoting angiogenesis and immune evasion (Schmitt et al. 2022).

The regulation and composition of the SASP have largely been investigated in model systems in which senescence is induced by DNA damage or genome instability (Rodier et al. 2009; Malaquin et al. 2016). Such triggers have been shown to induce expression of inflammatory genes through stimulation of NF-κB, a master regulator of the SASP (Chien et al. 2011b); this is coordinated via the DNA damage response kinase, ATM (Miyamoto 2011; Bournique et al. 2025). While DNA damaging agents are frequently applied in cancer therapeutic approaches, targeted therapies like CDK4/6i can induce senescent-like states through non-DNA damage mechanisms. This begs the question, what drives the SASP when there is cellular senescence but not DNA damage? Prior studies of the response to CDK4/6i treatment have thus far offered conflicting results regarding the involvement of NF-κB (Goel et al. 2017; Guan et al. 2017; Wang et al. 2022; Gleason et al. 2024; Lee et al. 2024).

Here we sought to investigate: (1) does the presence (or absence) of DNA damage give rise to distinct therapy-induced senescent cell states and SASP profiles over time? and (2) what epigenetic changes dynamically regulate the SASP in cancer cells in response to therapeutics? We chose to address these questions systematically in a liposarcoma model, a rare type of malignant mesenchymal tumor with limited treatment options in advanced disease. Over 90% of liposarcoma tumors are characterized by co-amplification of *CDK4* and *MDM2* (Creytens et al. 2015), and previous work has shown that CDK4/6i can induce a full senescence program with irreversible arrest driven by *ANGPTL4*, correlating with patient outcomes and clinical efficacy (Kovatcheva et al. 2015; Dickson et al. 2016; Gleason et al. 2024). While traditional chemotherapeutics, such as doxorubicin, trigger senescence through DNA damage, CDK4/6i instead arrest cells in G1 by preventing the activity of E2F transcription factors in a DNA damage-independent mechanism (Guan et al. 2017; Kovatcheva et al. 2017; Gleason et al. 2024) (Fig. 1A). We performed an extended 28-day time course directly comparing these two distinct triggers of therapy-induced senescence. By profiling epigenomic changes across multiple time points, we captured the temporal evolution of senescence-associated gene regulatory programs, including features that would be missed in short-term studies.

**Fig. 1:**
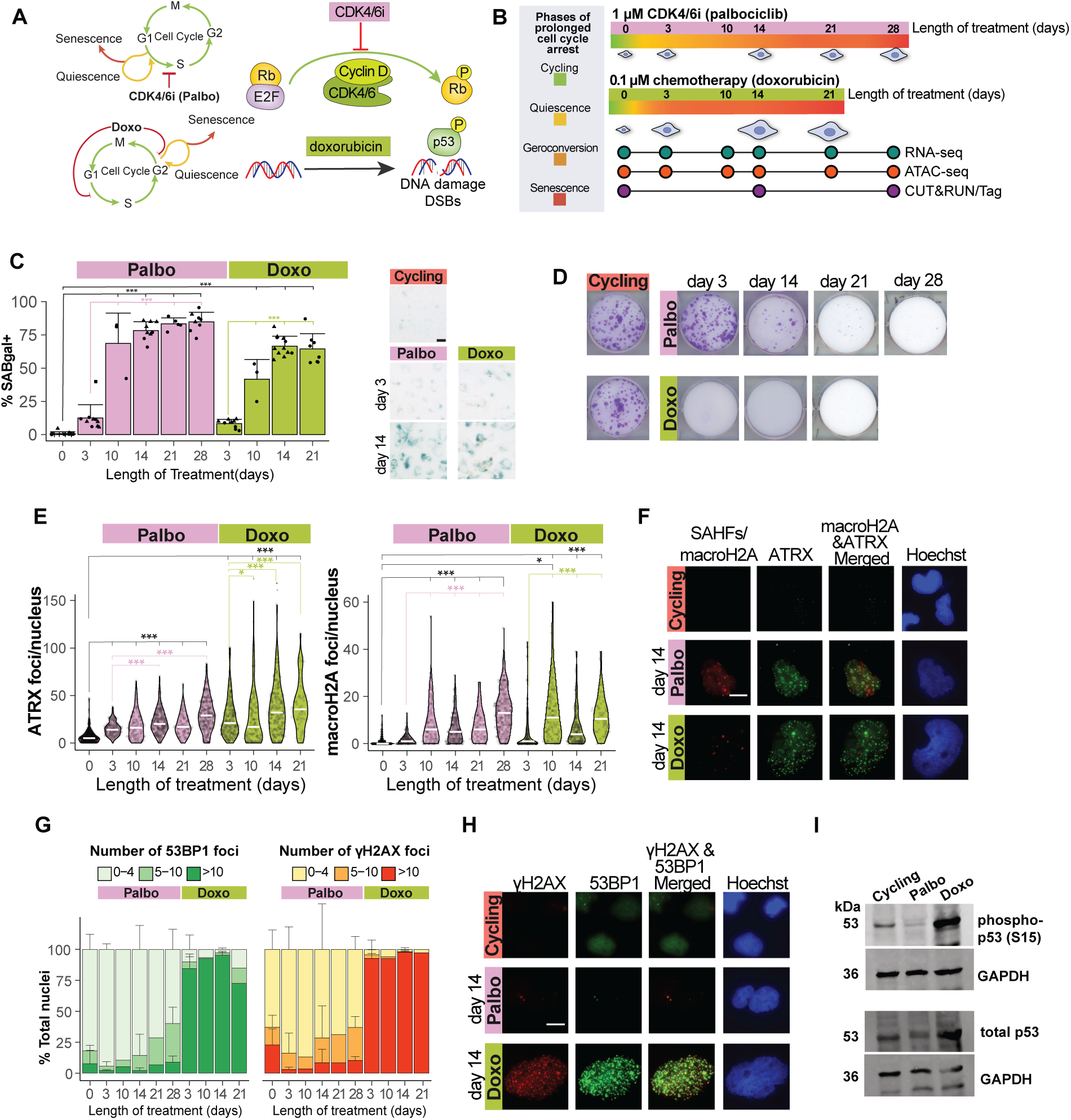
CDK4/6i and doxorubicin both induce senescence features with distinct levels of DNA damage and p53 activation. A. Diagram summarizing CDK4/6i (Palbo) vs. doxorubicin (Doxo) mechanisms of cell cycle arrest. B. Schematic overview of comparative time-course between CDK4/6i and doxorubicin and multi-omics approaches used to profile the transcriptome (RNA-seq), chromatin accessibility (ATAC-seq) and epigenome (CUT&RUN/Tag) in LS8817 cells. At least 2 biological replicates per timepoint and drug treatment were profiled. C. Left: Percent nuclei counted that had cellular SA-βgal+ staining at different timepoints. Number of nuclei counted was > 300 per condition at 10x objective magnification. Each point represents a field of view, with different shapes indicating biological replicates. Error bars denote standard deviation. Right: Representative SA-βgal staining images at day 3 and day 14 shown at 10x objective magnification with scale bar = 50 µm. D. Crystal violet staining after clonogenic outgrowth for 10-12 days. 800 cells were seeded per well in a 6-well plate in drug-free media after completion of drug treatment at each timepoint. E. Number of ATRX foci (left) and macroH2A foci (indicative of SAHFs; right) per nucleus was counted for ∼100 nuclei per timepoint. F. Representative ATRX and SAHF images are shown at day 14 of each drug treatment at 60x objective magnification. The scale bar is 15 µm. G. Number of γH2AX foci and 53BP1 foci per nucleus was counted for ∼100 nuclei per timepoint and stratified into 0-4, 5-10 or >10 foci/nuclei. H. Representative images of γH2AX and 53BP1 staining are shown at day 14 of each drug treatment at 60x objective magnification. The scale bar is 15 µm. I. Western blots of Serine 15-phosphorylated p53 (phospho-p53) and total p53 in Cycling, Palbo- and Doxo-treated LS8817 cell lysates harvested 18 days after treatment started. GAPDH was used as a loading control. Samples are from a separate experiment, independent of multi-omics data.

We find that early senescence programs are trigger-specific and largely remain distinct, including p53 activation following doxorubicin-induced DNA damage and an extracellular matrix (ECM)-enriched SASP program upon CDK4/6i treatment. At later time points, both therapies initiate a shared NF-κB-driven SASP, albeit with slower induction in the absence of DNA damage. Such activation of NF-κB is conserved beyond liposarcoma, observing a similar response in breast cancer cells. By extending the duration of CDK4/6i treatment here, we revealed delayed NF-κB activation and clarify its previously debated role from short-term studies, where its activity was not detected. Notably, NF-κB inhibition suppresses the shared SASP without reversing stable arrest, suggesting that selective modulation of SASP composition is possible while preserving growth arrest, representing new therapeutic opportunities to explore.

## RESULTS

### CDK4/6i and doxorubicin both induce senescence features with distinct levels of DNA damage and p53 activation

To understand whether the absence or presence of DNA damage drives distinct cell states over time, we compared 1 µM palbociclib (Palbo), a type of CDK4/6i, against 100 nM doxorubicin (Doxo), standard chemotherapy, over a 28-day period across multiple timepoints in a patient-derived dedifferentiated liposarcoma cell line, LS8817 (Singer et al. 2007; Brill et al. 2010; Crago and Singer 2011) (Fig. 1B). Among multiple well-differentiated/dedifferentiated liposarcoma (WD/DDLS) cell lines tested, LS8817 was among those which undergoes stable senescence in response to palbociclib (Kovatcheva et al. 2015; Kovatcheva et al. 2017), providing a unique and experimentally tractable system for interrogating CDK4/6i-induced senescence in a clinically relevant cancer type. To investigate the full breadth of gene regulatory networks that underlie the senescent state, we performed a time-resolved, multi-omics analysis that profiles the transcriptome using mRNA-seq, accessible chromatin regions by ATAC-seq, and the epigenome using CUT&Tag and CUT&RUN methods (Fig. 1B; Supplemental Table 1).

We first validated that cells are entering into a senescent state in response to both drug treatments. Since there is no universal biomarker of senescence, we measured several accepted hallmarks of senescence. We observed an increased amount of lysosomal activity, through senescence-associated beta galactosidase (SA-βgal) staining, in both CDK4/6i and doxorubicin over time (Fig. 1C). We confirmed cells entered a stable growth arrest state by assessing for clonogenic outgrowth upon drug removal (Fig. 1D). In these clonogenic outgrowth assays, doxorubicin caused a stable arrest in nearly the entire population early in treatment, by day 3, whereas stable arrest did not occur for most cells in response to palbociclib until day 14 of treatment (Fig. 1D). We also measured senescence-associated nuclear changes by immunofluorescence staining for the heterochromatin remodeler, ATRX (Kovatcheva et al. 2015; Kovatcheva et al. 2017) and the histone variant, macroH2A, a key structural SAHF component (Zhang et al. 2005) (Fig. 1E, F). Indeed, we saw that both treatments lead to an accumulation of both ATRX foci and SAHFs (Fig. 1E,F), indicating that the global chromatin landscape does undergo dynamic changes during senescence.

Given the differences in the mechanisms of action for each drug, we anticipated the amount of DNA damage would be a distinctive feature between treatments. We measured the foci of γH2AX, a classical marker of DNA damage, and 53BP1, a DNA repair factor (Fig. 1G,H). We found that even by day 28 of palbociclib treatment, the amount of DNA damage-associated foci remains comparable to that of untreated cycling controls (Fig. 1G). This contrasts with doxorubicin treatment, under which DNA damage is robustly induced and remains elevated across all timepoints (Fig. 1G). Next, we asked whether these differences in DNA damage are also reflected in the levels of phosphorylated p53, indicative of activation of the downstream DNA damage response (Sakaguchi et al. 1998; Saito et al. 2002; Loughery et al. 2014). Immunoblotting for phosphorylated p53 (Serine 15) and total p53 protein levels confirmed that CDK4/6i treatment does not result in downstream activation of p53, whereas p53 is robustly activated in doxorubicin (Fig. 1I). Thus, our results demonstrate that CDK4/6i treatment can induce senescence without the requirement for persistent DNA damage or activation of the DNA damage response.

### Global changes in the epigenome and transcriptome define drug-induced senescence trajectories

Given that both treatments drive cells into a senescent state, we asked whether the overall transcriptional regulatory state of these cells depends on the driver, particularly since doxorubicin triggers DNA damage while palbociclib does not. We assayed the regulatory landscape with ATAC-seq to measure chromatin accessibility. Sample distance measures and principal component analysis showed that the global transcriptome and chromatin accessibility landscape continues to change throughout senescence (Supplemental Fig. 1A). These overall changes correspond to the timing of the stable growth arrest state as observed by accumulation of SA-βgal and reduced clonogenic outgrowth (Fig. 1C,D).

We found that many changes in chromatin accessibility trended in the same direction, but the timing and magnitude of the change was often dependent on the treatment (Fig. 2A). A common feature among the senescent cells was decreased accessibility proximal to promoters (Fig. 2B: top). These regions are enriched in E2F motifs (Supplemental Fig. 1C,D), and thus their repression is indicative of cells disengaged from active cycling. Increases in accessibility, on the other hand, were predominantly localized to intergenic and intronic regions (Fig. 2B: bottom), suggesting potential enhancer activation. These regions are enriched for motifs of stress-responsive and lineage-specifying transcription factors (Balamurugan and Sterneck 2013; Kamachi and Kondoh 2013; Sarkar and Hochedlinger 2013; Golson and Kaestner 2016; Tremblay et al. 2018; Huh et al. 2019; Romano and Miccio 2020; Awasthi et al. 2021; Currey et al. 2021; Li et al. 2023; Ren et al. 2023) (Supplemental Fig. 1E), consistent with activation of transcriptional programs that shape senescence-associated cell fate decisions beyond cell cycle arrest.

**Fig. 2:**
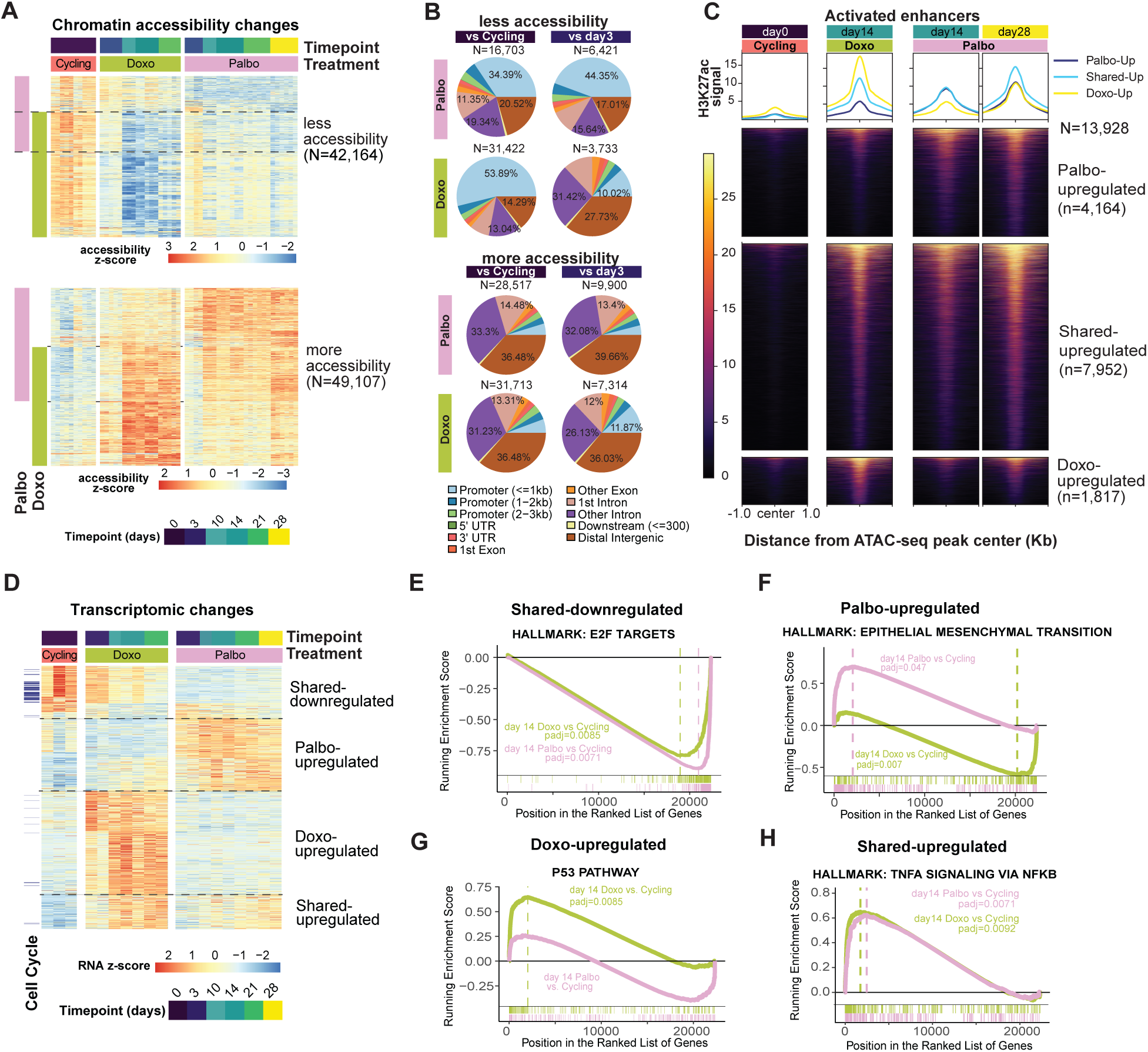
Distinct and shared transcriptional and chromatin responses define drug-specific senescence trajectories. A. Heatmap of z-scores from VST-normalized read counts under ATAC-seq peaks with significantly less accessibility (top) or more accessibility (bottom) over time, relative to cycling or day 3. Annotations indicate significance in Palbo (pink) or Doxo (green) and their genomic location. B. Genomic feature annotation of ATAC-seq peaks with less accessibility (top) or more accessibility (bottom), merged across all time points compared to Cycling (left) or day 3 (right)). N= number of peaks in merged peak-set making up the pie chart. C. RPGC-normalized H3K27ac coverage in drug-activated enhancers (total number N= 13,928). Each row in the heatmap represents a drug-activated enhancer: defined as a region of increased chromatin accessibility overlapping a corresponding H3K27ac region increasing in palbociclib (Palbo-upregulated), both treatments (Shared-upregulated) or doxorubicin (Doxo-upregulated) (See Supplemental Materials). The number of regions plotted is shown on the right. Scale bar represents bigwig signal of normalized coverage. D. K-means clustered heatmap of z-scores from rlog normalized counts for significantly changing genes across treatment and timepoints (days of treatment). Cell cycle genes annotated in dark green and blue, respectively. E. GSEA running enrichment score for enriched Hallmark term “E2F TARGETS” identified in Shared-downregulated genes. Ranked gene list is based on day 14 Doxo and day 14 Palbo relative to Cycling comparisons. *Padj* values showing significance of enrichment are shown. F. GSEA running enrichment score for enriched Hallmark term “EPITHELIAL MESENCHYMAL TRANSITION” identified in Palbo-regulated genes. G. GSEA running enrichment score for enriched Hallmark term “P53 PATHWAY” identified in Doxo-regulated genes. H. GSEA running enrichment score for enriched Hallmark term “TNFA SIGNALING VIA NFKB identified in Shared-upregulated genes. *Padj* values showing significance of enrichment are shown.

We wondered whether enhancer activation dynamics differ depending on the timing of senescence entry; to capture these changes, we profiled H3K27ac, a histone mark of active enhancers. We found that 28% of all peaks that increased in accessibility also displayed significantly increased H3K27ac signal (Supplemental Fig. 1F), indicating remodeling of the enhancer landscape underlying the therapy-induced senescent state. Accordingly, we defined these regions as drug-activated enhancers (Fig. 2C). Somewhat surprisingly, the majority of these activated enhancer elements were shared across treatment, albeit with different dynamics (Fig. 2C). H3K27ac in shared-upregulated enhancers gradually increases with CDK4/6i treatment, reaching levels similar to that of day 14 doxorubicin by day 28 (Fig. 2C). This is supported by principal component analysis, which shows that day 28 palbociclib clusters more closely with day 14 doxorubicin than day 14 palbociclib (Supplemental Fig. 1G). This delayed enhancer activation in the CDK4/6i treated cells corresponds to slower progression into stable growth arrest (Fig. 1D).

We next asked how the transcriptional landscape compared between these two distinct therapy-induced senescence states, across multiple time points (Fig. 1B; Fig. 2D; Supplemental Data 1). Consistent with decreased accessibility at promoter regions enriched in E2F motifs (Fig. 2B: top; Supplemental Fig. 1C,D), E2F target genes were downregulated in both drug treatments, reflecting the cell cycle arrested state (Fig. 2D,E; Supplemental Data 2). We also observed more obvious drug-specific and temporal effects than at the chromatin accessibility and enhancer activation level (Fig. 2D). This could be explained by differential transcription factor activity at DNA regulatory elements that are dependent on the DNA damage context. Among the genes that displayed higher expression levels in a drug-specific manner, palbociclib was related to a stronger “Epithelial to Mesenchymal Transition” signature (Fig. 2F; Supplemental Data 2). Phosphorylation of p53 at serine 15 has been shown to enhance p53 binding to chromatin, thereby facilitating transcriptional activation at p53-responsive promoters (Loughery et al. 2014). Supporting this notion, doxorubicin preferentially activated the p53 axis, as evidenced by induction of p53 target genes including CDKN1A (encoding p21) (Supplemental Fig. 1H,I), enrichment of the p53 pathway (Fig. 2G; Supplemental Data 2) that was absent in palbociclib, and increased accessibility at p53 motifs (Supplemental Fig. 1G).

Further analysis of drug-specific clusters revealed distinct early and late transcriptional responses within each treatment. Notably, while some late-response genes were also upregulated under the alternate condition, their induction was consistently greater in the treatment in which they were classified (Supplemental Fig. 2B: top). The early doxorubicin response was once again centered around p53 (Supplemental Fig. 2B: bottom; Supplemental Data 3), while the later response may involve metabolic adaptation through genes like *PPARGC1A* and *SORL1* (Supplemental Fig. 2B: bottom; Supplemental Data 3).

Although there was a large number of transcriptomic changes that were specific to the senescence trigger, we noted a set of genes that increased in expression as cells entered a senescent state independent of treatment (Fig. 2D: Shared-upregulated). This upregulation was faster in the doxorubicin treatment and more gradual with palbociclib, coinciding with the dynamics of the enhancer activation we noted previously (Fig. 2C). Gene Set Enrichment Analysis (GSEA) revealed this set of genes was enriched for NF-κB transcriptional activity (Fig. 2H; Supplemental Data 2), a motif we also observed among ATAC-seq peaks with shared increased accessibility between treatments (Supplemental Fig. 1E: middle; Supplemental Data 4).

### NF-κB is a predicted transcriptional regulator of the shared SASP in liposarcoma

Although multiple features of therapy-induced senescence vary between doxorubicin and palbociclib treatment responses (e.g., DNA damage activation, p53 signaling, and a large fraction of RNA and chromatin accessibility changes), we were particularly interested in what aspects these two very different drivers of stable arrest have in common. To investigate this, we focused on shared-upregulated genes which had distinct temporal dynamics in expression levels across drug treatments (Fig. 2D), with a strong enrichment in the “TNFA SIGNALING VIA NFKB” Hallmark term (Fig. 2H; Supplemental Data 2).

Despite its activity being undetected in prior short-term CDK4/6i treatment studies, the NF-κB complex and its canonical subunit, RELA (p65 or TF65) emerged as top predicted upstream regulators (Fig. 3A; Supplemental Data 2). We identified 40 NF-κB targets within the shared-upregulated gene module and analyzed their RNA levels at days 3 and 14 of treatment (Fig. 3B). Approximately half of these genes were also defined as SASP across multiple databases (Fig. 3B: right; Supplemental Data 1). Both drugs showed a significant increase in expression over time, consistent with a progressive NF-κB response (Fig. 3B). However, this response was delayed in palbociclib-treated cells, with the median fold change at day 14 resembling that of doxorubicin at day 3 (Fig. 3B). Corresponding to the shared induction of NF-κB target genes, NF-κB family motifs were similarly enriched in regions with increased chromatin accessibility in both drugs (Supplemental Fig. 3A; Supplemental Data 4). To confirm this shared NF-κB signature predicted by multi-omics analysis, we performed immunofluorescence on p65 (Fig. 3C; Supplemental Fig. 3B). In both treatments, we observe increased p65 nuclear localization, with timing that correlated approximately with the onset of stable cell cycle arrest (Fig. 1D), i.e., steadily increasing up to day 14 in response to palbociclib and much earlier, by day 3, in doxorubicin (Supplemental Fig. 3B).

**Fig. 3:**
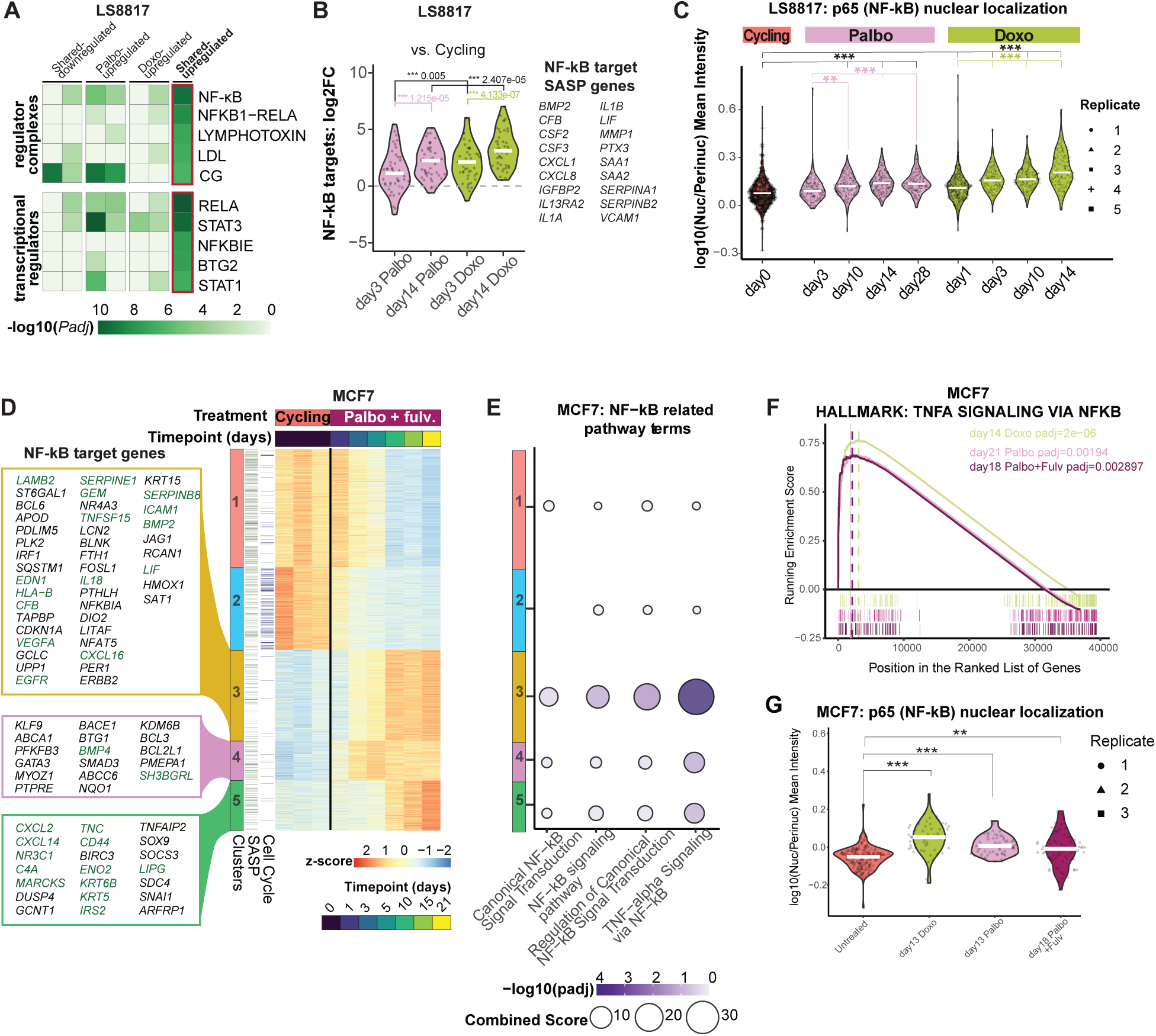
A shared senescence signature driven by NF-κB activation is delayed in CDK4/6i treatment across cancer cell types. A. -Log10(*Padj*) values of top 5 IPA predicted upstream regulator complexes and transcriptional regulators of Shared-upregulated genes relative to other RNA-seq clusters in LS8817 cells. B. Left: Log2 fold change (log2FC) values of NF-κB target genes that are part of the Shared-upregulated genes (RNA Cluster 7) in day 3 and day 14 Palbo and Doxo relative to Cycling in LS8817 cells. Median log2FC values for each condition are shown as a bold white line. The dotted gray line at log2FC = 0 indicates no change from Cycling. A paired Student’s t-test was used to assess significance between log2FC distributions. Right: gene IDs of NF-κB target genes that are also defined as SASP genes based on whether they were classified as such in manually curated databases provided in Wang et al.,(Wang et al. 2022) Gleason et al.,(Gleason et al. 2024) and the SASPAtlas (Basisty et al. 2020). C. Quantification of log10 ratio of mean intensity of p65 in the nucleus to perinuclear region in LS8817 cells. Statistical significance was calculated using Dunnett’s test. Each point is a cell that was quantified. D. K-means clustered heatmap of z-scores from rlog normalized counts for significantly changing genes across treatment and timepoints (days of treatment) based on RNA-seq performed in MCF-7 cells. SASP and cell cycle genes are annotated in dark green and blue, respectively. NF-kB target genes found in each upregulated gene cluster are shown left of the heatmap. Genes in dark green are also annotated as SASP. E. GO enrichment results of NF-kB related terms from each RNA-seq cluster in MCF7 cells. F. GSEA running enrichment score for enriched Hallmark term “TNFA SIGNALING VIA NFKB” for palbociclib, doxorubicin and palbociclib+fulvestrant treatment in MCF7 cells. *Padj* values from GSEA are shown. G. Quantification of log10 ratio of mean intensity of p65 in the nucleus to perinuclear region in MCF7 cells across different biological replicates. Statistical significance was calculated using Dunnett’s test.

### A shared NF-κB-driven SASP program is also found in MCF7 breast cancer cells

A previous study did not report NF-κB activation with 5 days of CDK4/6i treatment using the ER+ breast cancer model cell line, MCF7. Therefore, we wondered whether later timepoints would show NF-κB activation that could regulate the SASP in this system as well. To assess whether delayed NF-κB activation also occurred in this additional clinically relevant model, we treated MCF7 breast cancer cells with 1 µM palbociclib, 100 nM doxorubicin, or 100 nM palbociclib and 10 nM fulvestrant (Palbo+Fulv) —the standard of care regimen for ER+ breast cancer.

We confirmed that CDK4/6i induces senescence in MCF7 cells without triggering excess DNA damage, beyond what is observed in cycling cells, unlike doxorubicin (Supplemental Fig. 3C-E). We observed increased SA-βgal staining (Supplemental Fig. 3C) and reduced clonogenic outgrowth (Supplemental Fig. 3D) across all conditions.

While strong stable arrest is observed in Palbo+Fulv, few γH2AX and 53BP1 DNA damage-associated foci accumulated compared to doxorubicin (Supplemental Fig. 3E), consistent with what we see in the liposarcoma context (Fig. 1G,H).

Even in a model where NF-κB activity was previously undetected in short-term studies (Lee et al. 2024), we observed the eventual activation of NF-κB as a late treatment response. Time-resolved analysis over the course of 21 days of Palbo+Fulv treatment revealed 7,510 significantly altered genes based on a *Padj* threshold of < 0.05 (Fig. 3D). Strikingly, NF-κB-related pathways were enriched in a group of genes that are modestly upregulated at early time points (days 3-5) but show strong induction after day 10 (Fig. 3E, cluster 3). GSEA across different drug conditions in MCF7 samples, showed that the “TNFA SIGNALING VIA NFKB” Hallmark term was significantly enriched in both palbociclib- and doxorubicin-treated samples (Fig. 3F; Supplemental Data 2). Evidence of shared induction of NF-κB activity was further validated by immunofluorescence staining of p65, which showed increased nuclear localization across all drug treatments (Fig. 3G). This supports the notion that NF-κB-SASP network activation is a shared feature of therapy-induced senescence across cancer types.

To identify shared NF-κB–associated SASP components, we examined NF-κB target genes upregulated across treatments in MCF7 cells and found that 41 of 116 (35%) were common to palbociclib, Palbo+Fulv, and doxorubicin (Supplemental Fig. 3F). We next assessed conserved SASP programs driven by CDK4/6i across cell types. Several common SASP genes exist, including established NF-κB targets such as CXCL2 and CXCL8, which were consistently upregulated following CDK4/6i treatment in both MCF7 and LS8817 cells (Supplemental Fig. 3G). Although not a direct NF-κB target, *ANGPTL4*, a SASP factor necessary for CDK4/6i-mediated stable arrest (Gleason et al. 2024), was induced in both cell types. Moreover, *ANGPTL4* activates *IL1A*, a pro-inflammatory cytokine transcribed by NF-κB (Makulyte et al. 2026)—highlighting the complexity of the CDK4/6i-driven SASP, which may include components that can act either upstream or downstream of NF-κB. Together, these findings highlight a shared NF-κB response across distinct senescence-inducing treatments, alongside a CDK4/6i-specific SASP conserved across cancer cell types

### Nearby enhancer activation contributes to the induction of shared NF-κB target genes

Although NF-κB family motifs were globally enriched in regions of increased accessibility (Supplemental Fig. 3A), we sought to determine whether this enrichment was retained at regulatory elements more likely to directly influence expression levels of shared-upregulated genes. To focus on such elements, we restricted our analysis to nearby upregulated accessible regions, which were defined as within 50kb of genes belonging to the shared-upregulated gene module (Supplemental Data 5). Even under this constraint, we observed that increased Tn5 insertion coverage at TF65 (p65) motifs occurs across both drug treatments (Supplemental Fig. 4A-B).

Next, we identified nearby drug-activated enhancers by intersecting upregulated accessible regions with sites of increased H3K27ac, specifically located within 50kb of shared-upregulated genes that increased in expression (Fig. 4A). Notably, H3K27ac accumulation at these nearby enhancers was delayed in CDK4/6i relative to doxorubicin (Fig. 4A: top). Coinciding with the slower kinetics of NF-κB target gene induction in CDK4/6i (Fig. 2D; Fig. 3D), these shared-upregulated enhancers proximal to shared-upregulated genes were highly enriched in NF-κB family motifs, including canonical subunits TF65 and NFKB1 (Fig. 4B: left).

**Fig. 4:**
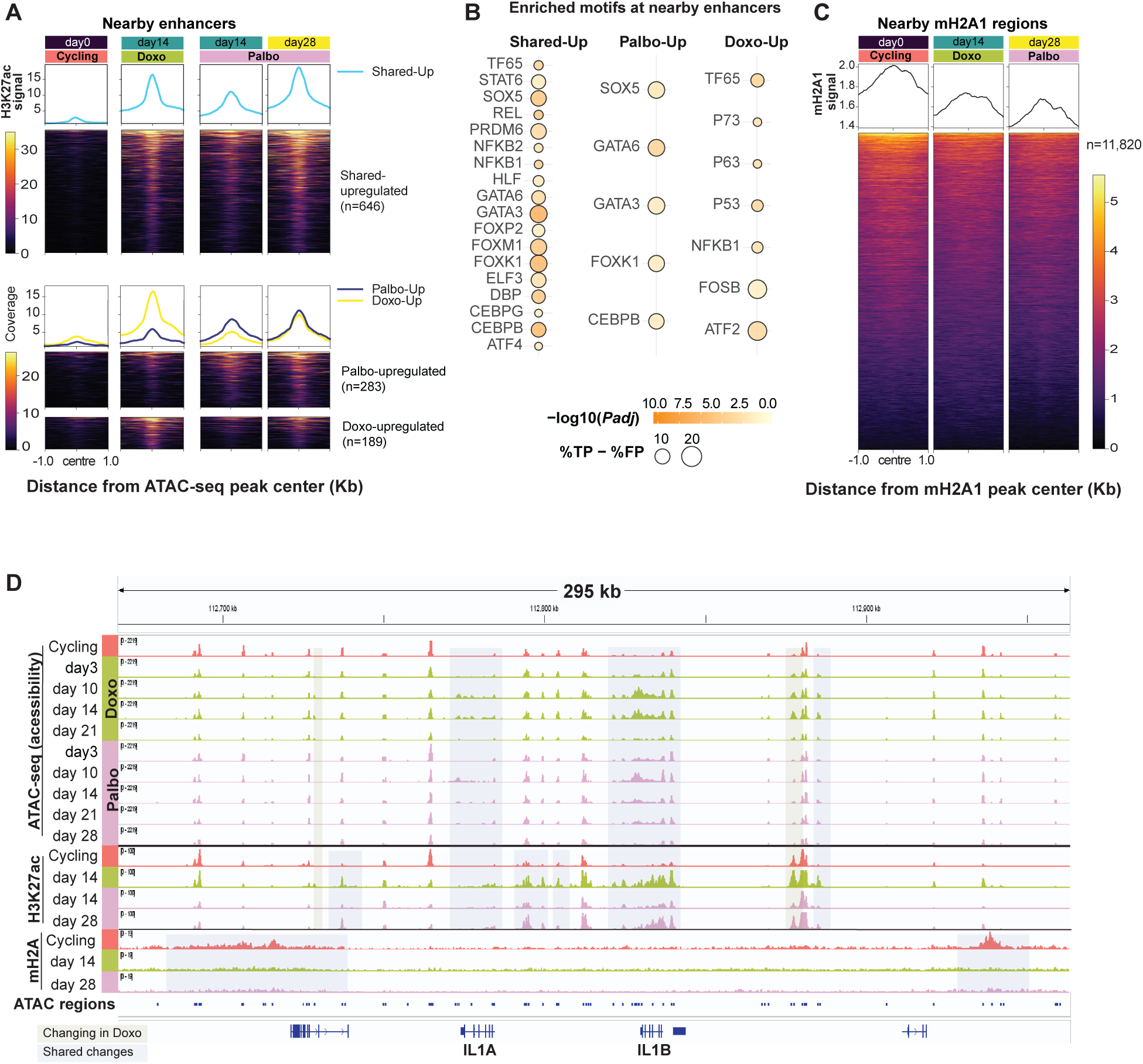
Activation and loss of macroH2A at enhancers near NF-κB-driven shared SASP genes. A. H3K27ac coverage signal across conditions in drug-activated enhancers within 50kb of shared-upregulated genes (Nearby enhancers). Each row in the heatmap represents a drug-activated enhancer: defined as a region of increased chromatin accessibility overlapping a corresponding H3K27ac region increasing in palbociclib (Palbo-upregulated), both treatments (Shared-upregulated) or doxorubicin (Doxo-upregulated) (See Supplemental Materials). The number of nearby peaks is shown on the left. Scale bar represents bigwig signal of normalized coverage. B. Motif enrichment results of drug activated enhancers shown in (A) within 50kb of Shared-upregulated genes. Dot intensity represents -log10(p-adj) values, while dot size reflects the difference between true positive and false positive peaks (%TP - %FP). C. macroH2A1 (mH2A1) signal for all mH2A1 peaks within 50kb of shared-upregulated genes (nearby mH2A1 regions) from Fig. 2D. The number of nearby mH2A1 peaks is shown on the left. Scale bar represents bigwig signal of normalized coverage. D. Genome browser view of *IL1A* and *IL1B* loci. Normalized bigwig tracks for ATAC-seq, H3K27ac and macroH2A1 (mH2A) are shown. Upregulated regions occurring in both treatments are highlighted in blue (Shared changes) while Doxo-upregulated regions are highlighted in green (Changes in Doxo).

In addition, motifs enriched in nearby palbociclib-upregulated enhancers—including CEBPB, GATA3, GATA6, SOX5, and FOXK1 (Fig. 4B: middle)—were also present at nearby shared-upregulated enhancers (Fig. 4B: top), suggesting continual engagement of transcription factors associated with mesenchymal enhancer activity and epithelial-mesenchymal transition programs. In contrast, doxorubicin-upregulated enhancers near shared-upregulated genes were additionally enriched for p53 family motifs (Fig. 4B: right), consistent with activation of the DNA damage response.

Interestingly, AP-1 motifs (e.g., FOSB and ATF2), a family of stress-responsive pioneer transcription factors (Biddie et al. 2011; Vierbuchen et al. 2017; Patrick et al. 2024) were also specifically enriched in doxorubicin-responsive enhancers near shared-upregulated genes (Fig. 4B: right). When considering regions gaining accessibility near the same genes, without requiring a change in acetylation, we found that there was some gain of accessibility at AP-1 motifs under palbociclib, but it was more modest than with doxorubicin (Supplemental Fig. 4C, top). Consistently, enrichment of AP-1 motifs in this accessible region set was also present at a low level (Supplemental Fig. 4C, bottom; Supplemental Data 5). However, when considering all upregulated ATAC-seq peaks genome-wide, peaks containing AP-1 motifs were strongly enriched at all times under doxorubicin treatment but also significantly enriched under palbociclib treatment, especially after day 3—exhibiting delayed activation kinetics (Supplemental Fig. 4D; Supplemental Data 4). Given that AP-1 has been identified as a master regulator of oncogene-induced senescence (Martínez-Zamudio et al. 2020), a DNA damage-driven process, the combined enrichment of p53 and AP-1 motifs at doxorubicin-upregulated enhancers may contribute to the earlier and more robust activation of NF-κB compared to CDK4/6i.

### Loss of macroH2A is associated with increased expression of NF-κB-driven shared SASP genes

In addition to enhancer activation, we wondered if other nearby changes in the epigenome contributed to activation of NF-κB-driven SASP genes. It is thought that the spatial rearrangement of heterochromatin into SAHFs plays a role in silencing cell cycle progression genes (Narita et al. 2003), while allowing for SASP genes to be sequestered into a transcriptionally permissive environment (Aird et al. 2016). Given the role that the histone variant, macroH2A, plays in transcriptional repression and the formation of the SAHFs (Zhang et al. 2005; Zhang et al. 2007), we used CUT&RUN to map the genome-wide distribution of macroH2A1 in late-stage therapy-induced senescence. We find that macroH2A2 is not expressed in this cell line (Supplemental Fig. 4E: top), additionally neither macroH2A1 RNA levels (Supplemental Fig. 4E: bottom) nor its global distribution on chromatin changed with drug treatment (Supplemental Fig. 4F). However, by examining macroH2A1-containing regions found within 50kb of shared-upregulated genes, we observe a striking loss of macroH2A (Fig. 4C). Hence, in addition to enhancer activation, locally reduced levels of macroH2A may also contribute to the activation of inflammatory SASP genes, likely through a de-repression mechanism.

We also focused on chromatin changes surrounding specific canonical SASP genes upregulated across both treatments. We examined a ∼300 kb window around the *IL1A, IL1B* (Fig. 4D), *CXCL1*, and *CXCL8* loci (Supplemental Fig. 4G). Across both treatments, we observed shared increases in chromatin accessibility as early as day 3, spanning gene bodies, transcription termination sites, and distal intergenic regions (Fig. 4D; Supplemental Fig. 4G). Despite these early accessibility gains, H3K27ac accumulation was delayed, remaining lower at day 14 in CDK4/6i-treated cells compared to doxorubicin before gradually increasing to comparable levels by day 28 (Fig. 4D; Supplemental Fig. 4G).

Finally, we identified enhancers selectively activated in doxorubicin-treated cells, which may contribute to the earlier and more robust induction of shared SASP genes observed under this condition (Fig. 4D; Supplemental Fig. 4G). Compared to ATAC-seq and H3K27ac signal, macroH2A1 coverage is broader and more diffuse, with signal depletion generally occurring in regions flanking—rather than within—the gene bodies of these inflammatory loci (Fig. 4D; Supplemental Fig. 4G). This pattern is surprising, and suggests an additional regulatory layer beyond the transcribed gene-body macroH2A depletion predicted by the “transcriptional pruning” model (Sun et al. 2018).

### The NF-κB-driven shared SASP component can be uncoupled from stable cell cycle arrest

We next considered whether NF-κB activation was necessary for cells to drive stable arrest, or is it solely responsible for the shared SASP program without any additional effects on senescence? We attempted to alter the NF-κB-driven shared SASP response through pharmacological inhibition with BAY11-7082 (BAY) (Pierce et al. 1997) during both treatments, after senescence was established (Fig. 5A). Principal component analysis shows NF-κB inhibition had minimal impact on the overall transcriptome, as BAY- and DMSO-treated samples clustered together within each drug group (Fig. 5B). However, BAY suppressed expression of key shared NF-κB target SASP genes, *CXCL1, CXCL8, IL1A, IL1B, GMCSF* (*CSF2*), *IL6* and *SOD2* (Fig. 5C; Supplemental Fig. 5A,B).

**Fig. 5:**
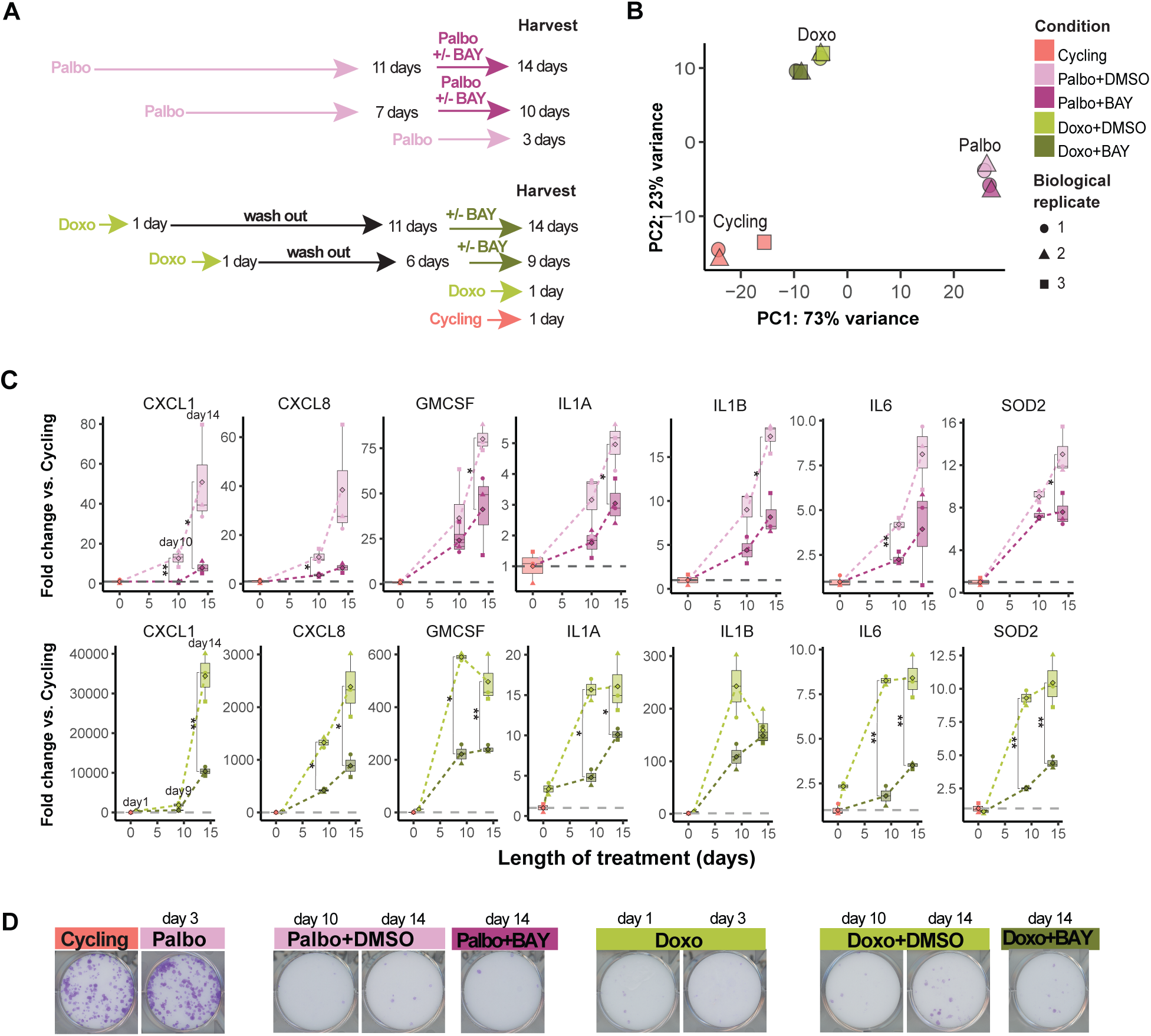
Inhibition of NF-kB activity suppresses the shared SASP without reversing stable growth arrest. A. Schematic of pharmacological inhibition of NF-κB with BAY11-7082 (BAY) in LS8817 cells (see Methods). 10 µM BAY or equivalent volume of DMSO (negative control) was added to cells for 2-3 days prior to harvest. B. Principal component analysis on RNA-seq of day14 palbociclib or doxorubicin treated LS8817 cells with or without NF-κB inhibitor, BAY11-7082 (BAY). C. RT-qPCR results of palbociclib or doxorubicin treated LS8817 cells with or without BAY. Relative fold change values to Cycling are shown. Dotted grey line indicates no change. Diamonds inside boxplot represent the mean fold change value. Circles, triangles, and rectangles represent the fold change of individual biological replicate. Dark purple and dark green shades indicate Palbo and Doxo samples treated with BAY while lighter purple and green shades indicate samples that had DMSO added at the same time as BAY samples. Statistical significance was calculated using a two-tailed t-test. D. Crystal violet staining after clonogenic outgrowth after palbociclib or doxorubicin treatment +/- BAY. 800 cells were seeded per well in a 6-well plate and grown in drug-free media for 12 days after harvest.

Beyond these, NF-κB inhibition downregulated 57 shared genes, as well as additional drug-specific targets (Supplemental Fig. 5C), suggesting that while NF-κB is a common feature of senescence, its ability to regulate specific targets may differ across treatments. Like previous studies of NF-κB in other types of senescence (Chien et al. 2011a; Lau and David 2019), inhibiting its activity was not enough to reverse senescence, as indicated by no differences in clonogenic outgrowth assays (Fig. 5D). Hence, although NF-κB activation does not appear to be necessary for maintaining stable arrest, it is responsible for driving part of the inflammatory SASP response.

### DNA damage response pathway is upstream of NF-**κ**B nuclear localization in doxorubicin but not CDK4/6i

NF-κB activity can be triggered by a variety of cellular stressors, among which the DNA damage response (DDR) is a major contributor (Miyamoto 2011). Notably, several contexts, such as mitochondrial dysfunction and treatment with histone deacetylase inhibitors, have demonstrated that senescence-associated DDR signaling can be engaged in the absence of overt DNA damage (Pospelova et al. 2009; Pazolli et al. 2012; Wiley et al. 2016).

While CDK4/6 inhibition does not directly induce DNA damage, these previous studies raise the possibility that DDR components could nonetheless be engaged through other mechanisms. To test whether DDR signaling contributes to NF-κB activation in this context, we pharmacologically inhibited the key DDR kinases ATM and ATR in arrested LS8817 cells (Supplemental Fig. 6A). We confirmed that the DDR signaling was dampened by ATM and ATR inhibition by immunoblotting for phosphorylated (phospho-) p53 (Fig. 6A; Supplemental Fig. 6B) and Chk2 (Fig. 6C; Supplemental Fig. 6C). Phospho-p53 is largely absent following palbociclib treatment, and co-treatment with ATM and ATR inhibitors did not alter phospho-p53 levels (Fig. 6A). Phospho-Chk2 levels were modestly increased under palbociclib treatment, suggesting mild ATM activation in the absence of p53 activation and additional DNA damage (Fig. 6B). In contrast, doxorubicin treatment induced robust p53 and Chk2 phosphorylation and activation, which was attenuated by ATM and ATR inhibition (Fig. 6A,B). This corresponded with reduced p65 nuclear localization, consistent with DNA damage-dependent activation of NF-κB (Fig. 6C). In contrast, inhibiting ATM and ATR did not affect NF-κB nuclear localization in CDK4/6i-treated cells (Fig. 6C), indicating that NF-κB activation in this setting occurs independently of the canonical DDR pathway.

**Fig. 6:**
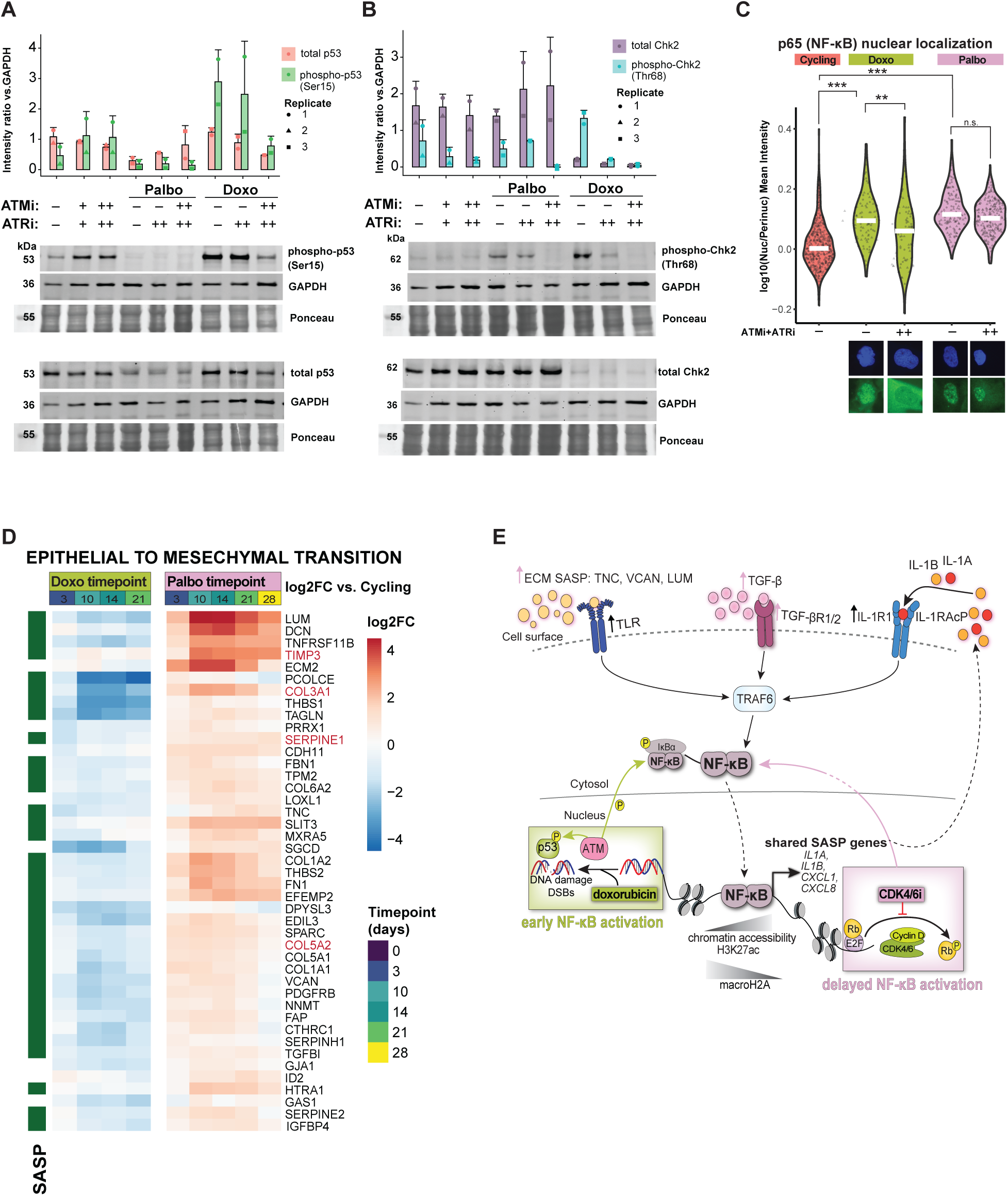
Inhibiting ATM and ATR affects NF-κB nuclear localization in doxorubicin but not CDK4/6i therapy-induced senescence. A. Western blots of LS8817 cells treated with Palbo (17 days) or Doxo (7 days), with or without ATR inhibition (ATRi; 4 μM AZD6783) or combined ATR and ATM inhibition (ATRi + ATMi: 4 μM AZD6783 + 4 μM KU-60019) for two biological replicates (Bioreps). ATR and ATM inhibitors were added after cell-cycle arrest had been established. ImageJ quantification of signal relative to GADPH loading control (Top). Phospho-p53 (Ser15) relative to total p53 (Bottom). “–” indicates DMSO control; “+” and “++” indicate 2 µM and 4 µM inhibitor, respectively. Replicate indicates different biological replicates. B. Same as (A) but western blots for phospho-Chk2 (Thr68) relative to total Chk2. C. Quantification of log10 ratio of mean intensity of p65 in the nucleus to perinuclear region in LS8817 cells treated with ATMi+ATRi or DMSO. Statistical significance was calculated using Dunnett’s test. Representative images taken at 60x objective magnification of different amounts of p65 nuclear localizations in each condition are shown at the bottom. D. Log2 fold change (log2FC) values of Palbo-upregulated genes that are under the Hallmark term “EPITHELIAL TO MESENCHYMAL TRANSITION”, across different timepoints (days of treatment) and drug conditions, relative to Cycling or day 3 controls for each drug treatment. Genes that are also SASP are shown on the left in dark green. Genes that are also NF-κB targets are shown in red text. E. Summary model showing possible pathways activated by CDK4/6i and doxorubicin to induce the shared SASP driven by NF-κB at different temporal rates.

### ECM driven SASP is specifically activated in CDK4/6i-induced senescence

Given that inhibition of DNA damage response factors did not affect NF-κB nuclear localization in CDK4/6i-treated cells, we next sought to identify alternative upstream pathways by characterizing gene expression programs specific to CDK4/6i-induced cell cycle arrest. Notably, genes associated with TGF-β signaling, including ligands (Supplemental Fig. 6D) and receptors (Supplemental Fig. 6E), were selectively upregulated in CDK4/6i-treated cells, but not in doxorubicin-treated cells. In addition, many of these genes encode ECM components and established TGF-β targets, several of which have been identified as SASP genes in other contexts (Fig. 6D). These findings suggest that, in parallel to the later NF-κB-driven shared SASP, CDK4/6 inhibition promotes an earlier, ECM-enriched SASP program.

Consistent with a role for extracellular signaling in this context, *IL1R1*, the primary receptor that *IL1A* and *IL1B* binds to (Wesche et al. 1997), increases in expression during CDK4/6i treatment (Supplemental Fig. 6F: top). This indicates potential activation of cell surface receptors that could drive downstream NF-κB activation. We also detected increased expression of TLR4 in both treatment conditions (Supplemental Fig. 6F: bottom), pointing to pattern recognition receptor signaling as an additional route to NF-κB activation (Kelsh and McKeown-Longo 2013; Guo et al. 2024). We hypothesize that upregulation of multiple cell surface receptors—including *TGFBR1/2, IL1R1*, and *TLR4* (Supplemental Fig. 6D-F)—along with ECM-enriched SASP components (Fig. 6D) that engage these receptors, may provide alternative routes to NF-κB activation in CDK4/6i-induced senescence, independent of intrinsic DNA damage signaling (Fig. 6E).

## DISCUSSION

Therapeutic response in various cancers is in part determined by how effectively tumors are driven into stable growth arrest. However, even upon entering such a senescent state, the profile of secreted molecules, or SASP, plays an important role in influencing the neighboring environment (Wang et al. 2024). Unlike replicative senescence that arises over a lifetime (McHugh and Gil 2018), therapy-induced senescence-like states are acute responses to cancer treatments, exhibiting many features of canonical senescence although mediated through distinct mechanistic triggers (Wagner and Gil 2020; Wang et al. 2020; Schmitt et al. 2022). Such acute models of senescence offer new insights into the underlying networks that are accessed during these processes and assist us in revealing common underlying features of the elusive core components of what it means to be a senescent cell. Additionally, understanding how the composition of the SASP changes over time across different cancer therapies will be essential in identifying new targeted approaches that improve patient outcomes.

Here we have used a dedifferentiated liposarcoma patient-derived cell line known to enter senescent-like states in response to CDK4/6i, and identified NF-κB as a key regulator of the common SASP inflammatory response. We show that this module of the response arises despite being triggered by a DNA-damaging agent (i.e., doxorubicin) or a DNA damage-independent mechanism (i.e., the CDK4/6 inhibitor palbociclib), and can be decoupled from stable arrest. We confirm this shared NF-κB activation also occurs in an estrogen receptor-positive (ER+) breast cancer cell line model (MCF7), supporting the generality of this regulatory node as a key feature of therapy-induced SASP. While the use of cell line models does not allow for the assessment of the SASP’s effects on immune clearance and/or evasion, we sought to focus on the molecular landscape of the tumor cell to clarify this context. Therefore, we mapped the temporal dynamics of epigenomic and transcriptomic networks at different phases of the CDK4/6i-induced senescence, by analyzing the genome-wide chromatin accessibility, H3K27ac-marked enhancers, and mRNA levels at early quiescent (day 3), mid transition (day 14), and late senescent (day 28) timepoints (Fig. 1B). We compared this to the DNA damage-induced senescent state that arises rapidly upon doxorubicin exposure.

Most transcriptomic changes are drug-distinct differences explained by p53 activation specific to DNA damage induced by doxorubicin (Fig. 2D,G) and a CDK4/6i-specific treatment response characterized by increased expression of ECM-related genes (Fig. 2D,F; Supplemental Fig. 2B). Nonetheless, we identified a common SASP-associated enhancer module enriched for NF-κB motifs that displayed delayed activation in CDK4/6i relative to doxorubicin, which was further substantiated by later accumulation of H3K27ac at enhancer elements (Figs. 2B,3A-B,4A-B). A concurrent loss of the SAHF component macroH2A around these loci may further contribute to the induction of the shared SASP module (Fig. 4C). While loss of macroH2A has been linked to de-repression of inflammatory loci in oncogene-induced senescence (Chen et al. 2015), to our knowledge this is the first report of its depletion in CDK4/6i-induced senescence, although its functional role here remains to be determined.

Our data reveal a previously underappreciated inflammatory component of CDK4/6i-induced senescence, with potential implications for long-term therapeutic outcomes. Although NF-κB’s role in senescence induced by oncogene activation is established (Chien et al. 2011a; Salminen et al. 2012), its activation in CDK4/6i-induced senescence was surprising, as prior studies failed to detect NF-κB activity (Goel et al. 2017; Wang et al. 2022; Lee et al. 2024). This discrepancy likely reflects differences in the duration of treatment, as these studies were focused on early responses. We did not detect NF-κB activation in earlier timepoints, in line with a previous observation in MCF7 cells after five days of treatment (Lee et al. 2024). However, by increasing the duration of CDK4/6i treatment (i.e., 14, 21, 28 days) we observed delayed but robust NF-κB activation across both liposarcoma and breast cancer cells.

To determine whether the NF-κB-driven inflammatory response is necessary for the cell to enter stable arrest, we showed that important shared SASP factors *(CXCL1, CXCL8*, *IL6*, *IL1A*, and *IL1B*) can be effectively suppressed by NF-κB inhibition *without reversing stable arrest* (Fig. 5 A-D). Many of these shared SASP factors have known tumor-promoting functions, including immunosuppression, metastasis, and chemoresistance (Ancrile et al. 2007; Acosta and Gil 2009; Jones and Jenkins 2018; Lau and David 2019; Rebe and Ghiringhelli 2020; Garlanda and Mantovani 2021; Wu et al. 2022). The SASP may therefore be tunable by adjuvant therapies to diminish pro-tumorigenic side effects.

Beyond shared targets, CDK4/6i also induces a distinct ECM-enriched SASP program (Fig. 6D) that may be driven by TGF-β signaling (Supplemental Fig. 2B,6D-E) and its downstream transcription factors (Fig. 4B). Future work is needed to understand the interplay and regulation between CDK4/6i-distinct and shared components of the SASP. Nonetheless, our data support a growing shift in the way we think about the clinical implications of CDK4/6i treatment. While it may offer initial benefits to patients over traditional chemotherapy, over time it leads to the emergence of a pro-inflammatory SASP driven by NF-κB that can promote tumor progression. Further therapeutic options that modulate the SASP while maintaining arrest are an area ripe for development.

These findings improve our understanding of the questions we set out to answer regarding the dependence of the SASP on DNA damage and the dynamic nature of the epigenetic changes that underlie the response. Since macro was previously shown to activate enhancers in response to CDK4/6i in breast cancer (Watt et al. 2021), we asked whether increased accessibility at AP-1 motifs contributed to the differences in the temporal activation of the SASP. In our data, both drug treatments showed evidence of AP-1 activation but exhibited distinct kinetics: rapidly increased in doxorubicin and more gradual with CDK4/6i (Supplemental Fig. 4C-D), with additional AP-1 enrichment at Doxo-specific increased H3K27ac regions. This may indicate that doxorubicin triggers full enhancer activation at AP-1 bound sites, whereas palbociclib treatment merely causes AP-1 factors to open chromatin via pioneering activity (Biddie et al. 2011; Vierbuchen et al. 2017). AP-1 is known to act synergistically with NF-κB to drive the expression of its target genes through genome-wide chromatin remodeling (Fujioka et al. 2004; Oeckinghaus et al. 2011; Ji et al. 2019), which offers an explanation for the timing and magnitude differences of NF-κB-driven SASP activation across treatments; earlier and robust in doxorubicin but delayed in CDK4/6i.

An outstanding question is, what drives NF-κB activation in CDK4/6i-induced senescence in the absence of DNA damage? Inhibition of ATM and ATR does not affect NF-κB nuclear localization with CDK4/6i but indeed blocks this response with doxorubicin (Fig. 6C), supporting a DNA damage response-independent mechanism in the CDK4/6i context. Therefore, NF-κB activation in CDK4/6i treatment is likely mediated through alternative upstream pathways, which we suspect includes TGF-β and IL-1, known mediators of this pathway (Guo et al. 2024). Both *TGFBR1/2* and *IL1R1* expression levels are upregulated in CDK4/6i treatment (Supplemental Fig. 6D-F), and converge on the TRAF6-TAK1 signaling axis, resulting in nuclear localization of NF-κB (Freudlsperger et al. 2013; Wang et al. 2021). In addition, both CDK4/6i and doxorubicin treatment cause increased expression of *TLR4*, another cell surface receptor that feeds into NF-κB activation (Guo et al. 2024). *TLR4* can be activated by endogenous damage-associated molecular patterns (DAMPs), including ECM components, versican (*VCAN*) and tenascin C (*TNC*) (Kelsh and McKeown-Longo 2013; Han et al. 2020), which are part of the CDK4/6i-enriched SASP (Fig. 6E). Activation of NF-κB promotes continued upregulation of *IL1A* and *IL1B* (Figs. 3B,4D), resulting in a positive feedback loop that sustains NF-κB activity (Croston et al. 1995; Weber et al. 2010). Together, our data suggest a model to be tested in future studies (Fig. 6E), whereby NF-κB activation in CDK4/6i-treated cells arises gradually through upregulation of cell-surface receptors and extracellular signaling rather than acute DNA damage.

Our results underscore two important considerations: (1) the importance of the underlying chromatin landscape that can vary intrinsically by cell type and extrinsically by trigger response, and (2) the dynamic and tunable nature of the SASP. In conclusion, we propose a model in which both doxorubicin and CDK4/6i induce senescence through distinct molecular mechanisms that converge over time, leading to NF-κB activation through enhancer remodeling and loss of macroH2A. These epigenetic changes promote a shared senescence response that manifests as a common set of cytokines and chemokines, which are secreted as part of the SASP. Our results bring new insights that help explain conflicting reports on various features of therapy-induced senescence, including the involvement of NF-κB in therapeutics that act via a DNA damage-independent mechanism.

## MATERIALS AND METHODS

### Cell lines and cell culture

Human LS8817^TetON^ ^FLAG-MDM2^ liposarcoma cell line was a gift from Dr. Andrew Koff’s laboratory at Memorial Sloan Kettering Cancer Center and has been extensively characterized in the literature (Singer et al. 2007; Barretina et al. 2010; Brill et al. 2010; Gobble et al. 2011; Miller et al. 2013; Kovatcheva et al. 2015). LS8817 have previously been described using the nomenclature DDLS8817 ([RRID:CVCL_M814](https://www.cellosaurus.org/CVCL_M814)). Cells were cultured in DMEM high glucose no glutamine (Cat# 11960044, Gibco), supplemented with 9% Fetal Bovine Serum (Cat# 10082147, Gibco™), 1% of 100x GlutaMAX™ Supplement (Cat# 35050061, Gibco™) and 1% of 10,000U/mL Penicillin/Streptomycin (Cat# 15140122, Gibco™) and maintained at 37°C and 5% CO_2_. Cells were authenticated by performing STR profiling to another LS8817 isogenic cell line. Cells were passaged at a 1:3 ratio every 3 days and regularly tested for mycoplasma.

Human MCF-7 cells (RRID:CVCL_0031) were cultured in in 75 cm2 flasks DMEM/F-12 (#Cat 10565018, Thermo Fisher Scientific) supplemented with GlutaMAX, 10% Fetal Bovine Serum (Cat# F4135, Sigma-Aldrich) and 1% Penicillin-Streptomycin at 37 °C and 5% CO_2_. Prior to their use, MCF7 cell identity was verified by STR profiling.

### Drug treatments

Cells were passaged at a 1:3 ratio one day before drug treatment. For samples sent for sequencing, cells were treated with either 1 µM palbociclib (Cat# S1116, Selleck Chemicals) or 100 nM doxorubicin (Cat# S1208, Selleck Chemicals), with media changes every 2–3 days until harvest.

For NF-κB inhibition experiments, 10 µM BAY11-7082 (Cat# S2913, Selleck Chemicals) was co-administered with 1 µM palbociclib in cells that had been treated with palbociclib 3 days before harvest. For the corresponding timepoints in Doxo, cells were treated with 100 nM doxorubicin for 24 hours, after which the drug was washed out. 10 µM BAY11-7082 was then added for 2–3 days prior to harvesting the Doxo-treated cells.

For ATMi + ATRi experiments, LS8817 cells were first treated for 3 days with 1 µM palbociclib and 100 nM doxorubicin. At day 3, cells treated with palbociclib continued to receive palbociclib with the addition of ATRi (AZD6783 at 4 µM; Cat #T3338, TargetMol), ATMi + ATRi (4 µM KU-60019; Cat #S1570, SelleckChem + 4 µM AZD6783) or equal volume of DMSO as vehicle control. Doxorubicin was washed out after ∼3 days before addition of DMSO, ATMi, ATRi or ATMi+ATRi treatment until harvest. Fresh drug-containing media was changed every 2-3 days until harvest.

Stock solutions were prepared as followed: palbociclib (10 mM in ultrapure dH₂O, stored at -80°C), doxorubicin (1 mM in ultrapure dH₂O, stored at -20°C), BAY11-7082 (50 mM in DMSO, stored at -80°C), KU-60019 (10 mM in DMSO,, stored at -80°C), AZD6783 (10 mM in DMSO, stored at -80°C)

### Immunofluorescence

50,000-60,000 cells were plated on acid washed glass coverslips and allowed to adhere overnight. Cells were fixed for 15 minutes with 4% formaldehyde followed by permeabilization with 0.1% Triton-X. Blocking was done with 0.5% Tween-20 and 1% normal goat serum for 30 minutes-1 hour at room temperature (RT).

Antibody dilutions used: ATRX (Cat# A301-045A, Bethyl) 1:2000, macroH2A (Cat# sc-377452, Santa Cruz) 1:500, γH2A.X (Cat# 05-636, Millipore) 1:1000, 53BP1 (Cat# ab172580, Abcam) 1:1000, NF-κB p65 (Cat #8242, Cell Signaling Technology) 1:800.

Primary antibody incubations were done at 4°C overnight. The next day, secondary Alexa Fluor™ antibodies (Cat #A-11004, A-11008, Invitrogen) were diluted 1:500 with 1:2000 Hoechst (Cat #H3570, Fisher Scientific) and incubated for 1 hour at RT. All reagents, including antibodies, were diluted in 1X PBS. All steps were followed by 3 washes of 1X PBS. Coverslips were mounted on homemade mounting medium (1 mL 10X PBS, 9 mL Ultrapure glycerol and 50mg of N-propyl gallate (Cat# A10877-06, Alfa Aesar)).

All images were acquired using the Nikon Ti2E Research Microscope System where about 100 nuclei or more were used for downstream quantification. To quantify ATRX, macroH2A, γH2AX, and 53BP1 foci in immunofluorescent images, we used a custom CellProfiler pipeline (Stirling et al. 2021). Foci and nuclei were identified using the **IdentifyPrimaryObjects** module, with the **EnhanceOrSuppressFeatures** module applied beforehand to improve foci detection. The **RelateObjects** module was then used to associate foci with their respective nuclei. The NF-κB nuclear-to-perinuclear intensity ratio was calculated using a CellProfiler pipeline based on https://www.youtube.com/watch?v=DlQSgDIGuds. Briefly, **IdentifyPrimaryObjects** detected nuclei, followed by **ExpandOrShrinkObjects** to expand nuclear boundaries, and **IdentifyTertiaryObjects** to define the perinuclear region. The **MeasureObjectIntensity** module was then used to quantify intensity in nuclear and perinuclear regions. SA-βgal+ associated nuclei were manually counted using Fiji software (Schindelin et al. 2012). For all quantifications, Dunnett’s test was used to test for significance between pairwise comparisons of conditions.

### SA-βgal staining

Senescence-associated beta galactosidase (SA-βgal) staining was conducted following the manufacturer’s protocol (Cat #9860S, Cell Signaling Technology), with all reagents diluted in 1× PBS instead of dH2O.

### Colony formation assays

Clonogenic outgrowth was performed in a 6-well plate by plating 1600, 800, and 400 cells per well using a 2× serial dilution method. Cells were allowed to grow for 10-12 days in drug-free media, followed by staining with crystal violet (1.25 g in 100 mL methanol and 400 mL distilled water) for 10 minutes at RT. Plates were scanned using an Epson Perfection V600 photo scanner.

### Western blots

Cells grown in a 10cm dish were washed once with ice-cold 1x Tris Buffer Saline (TBS), then scraped into 2mL of ice-cold 1x TBS contained 1x phosSTOP (Cat #04906845001, Millipore Sigma) & 1x cOmplete™ EDTA-free Protease Inhibitor (Cat #11836170001, Millipore Sigma). Cells were spun down at 400g for 10 minutes at 4oC, before removing the supernatant. 100-200 µL of 1x Laemmli buffer (Cat #1610747, Bio-Rad) + 50mM Dithiothreitol (DTT) + 10% glycerol + 1x phosSTOP & 1x cOmplete™ EDTA-free Protease Inhibitor was added to the cell pellet before boiling at 95°C for 5 minutes. Cell lysates were then sonicated for 3 cycles of 10 seconds on and 25 seconds off on the Bioruptor.

4–20% Mini-PROTEAN® TGX™ Precast Protein Gels (Cat # 4561096, Bio-Rad) were ran in 1x Tris/Glycine/SDS buffer (Cat #1610732) at 200V for 30 minutes then transferred to a nitrocellulose membrane (Cytiva, 0.1 µm pore size, Cat #10600000) overnight at 4ᵒC in 1x transfer buffer [20 mM Tris, 150 mM Glycine, 20 % Methanol] (14V). Membranes were blocked for 30 min at RT with Intercept® (TBS) Protein-Free Blocking Buffer (Cat# 927-80001, LICORbio), followed by primary antibody incubation (diluted in Intercept® T20 (TBS) Antibody diluent (Cat# 927-65001, LICORbio)) for overnight in 4°C. Membranes were washed then incubated 1:10,000 dilution with secondary antibodies (Cat #926-32211, Cat #926-68070, LICORbio) for 1 hour at RT and washed. Blots were then imaged on a LI-COR Odyssey M system. Each wash step occurred 3x, 5 min each wash, in 1x TBS + 0.05% Tween-20.

The following antibodies were used: Phospho-p53 (Ser15) Antibody (Cat #9284, Cell Signaling Technologies) 1:1000, p53 Antibody Cat #9282) 1:1000, GAPDH (D4C6R) Mouse Monoclonal Antibody, (Cat #97166, Cell Signaling Technologies), Phospho-Chk2 (Thr68) (C13C1) Rabbit Monoclonal Antibody (Cat #2197, Cell Signaling Technology) 1:1000, Chk2 (D9C6) Rabbit Monoclonal Antibody (Cat #6334, Cell Signaling Technology) 1:1000

Signal intensities of protein bands relative to loading control was quantified following the ImageJ protocol described by (Davarin 2015).

### RNA-sequencing

RNA extraction was done on at least 2 biological replicates per condition using the Quick-RNA Microprep Kit (Cat #R1050, Zymo Research). Sample quality was checked by determining RNA integrity number (RIN) via running samples through High Sensitivity RNA ScreenTape (Cat #5067-5579, Agilent Technologies) on the Tapestation.

LS8817 cells: Samples were sent to Novogene for stranded library preparation and sequencing. cDNA libraries were made with NEBNext® Ultra™ II Directional RNA Library Prep Kit for Illumina (Cat #E7765, New England Biolabs) and sequenced on Illumina NovaSeq 6000 Sequencing System as 150-base pair-end reads.

### ATAC-sequencing

ATAC libraries were generated following Corces et al. protocol for OMNI-ATAC (Corces et al. 2017). In brief, cells were treated in culture medium with DNase (Cat #LS002007, Worthington) at a final concentration of 200 U/mL at 37 °C for 30 minutes, then washed twice with PBS at 500 RCF, RT. Approximately ∼50,000-60,000 cells were used for each transposition reaction. Following permeabilization, 2.5 µL of Illumina TDE1 enzyme (Cat# 20034198, Illumina) was used for transposition at 37 °C for 30 minutes. Libraries were PCR amplified for 8-13 cycles with Illumina i5 or i7 barcoded primers and were cleaned up pre- and post-amplification following the manufacturer’s instructions using Zymo DNA Clean and Concentrator-5 Kit (Cat# D4014, Zymo Research). At least 2 technical replicates and 2 biological replicates per condition were sequenced. Fragment length distribution and DNA amount was quantified via TapeStation and Qubit to assess for quality prior to sequencing. Libraries were sequenced on the Illumina NovaSeq 6000 or on the NextSeq 1000 Sequencing System.

### CUT&Tag sequencing

CUT&Tag on LS8817 cells was performed following the protocol from Kaya-Okur et al. (Kaya-Okur et al. 2020) and was previously described in Soroczynski et al. (Soroczynski et al. 2024). Antibodies used include: Anti-Histone H3 (acetyl K27) antibody - ChIP Grade (ab4729, Abcam), 1:100; Guinea Pig anti-Rabbit IgG (Heavy & Light Chain) antibody (ABIN101961, Antibodies-online), 1:100.

### CUT&RUN sequencing

CUT&RUN in LS8817 cells was performed on macroH2A1 (Cat #ab37264, lot #1046297-1, Abcam), 1:100 following an optimized protocol by Epicypher ([Version 2.2](https://www.epicypher.com/wp-content/uploads/cutana-cut-and-run-protocol_v2.2.pdf)). NEBNext Ultra II DNA Library Prep Kit for Illumina was used for fragmentation, end repair and dA-tailing, adapter ligation and PCR amplification according to the manufacturer’s instructions. Ampure XP beads were used for post-PCR cleanup. Samples were eluted in IDTE buffer.

### RT-qPCR

Total RNA was extracted using the Quick-RNA Microprep Kit (Cat# R1050, Zymo Research) from at least two biological replicates per condition. cDNA was synthesized with the Applied Biosystems High-Capacity cDNA Reverse Transcription Kit (Cat# 4368814, Thermo Fisher Scientific) and SUPERase• In™ RNase Inhibitor (Cat# AM2696, Thermo Fisher Scientific). Prior to qPCR, cDNA was diluted to 5 ng/μL in dH2O.

qPCR was performed in triplicate in a 96-well microplate using a 15 μL reaction containing 7.5 μL PowerUp SYBR Green Master Mix (Cat# A25742, Thermo Fisher Scientific), 4 μL dH2O, 0.75 μL each of forward and reverse primers, and 1.5 μL of 5 ng/μL cDNA. Reactions were run on a QuantStudio™ 5 Real-Time PCR System with the following untreated cycling conditions: 98°C for 30 seconds, followed by 40–45 cycles of 98°C for 10 seconds, 63°C for 30 seconds, and 72°C for 1 minute. Cq values were normalized to the housekeeping gene *PUM1*, which was selected based on its stable expression across conditions in RNA-seq datasets.

All sequencing datasets have been deposited in the Gene Expression Omnibus (GEO) database. Code supporting analysis of sequencing datasets can be found at https://github.com/yeungj234/Manuscript_Figures. Further information and request for resources and reagents should be directed to and will be fulfilled by the lead contact, Dr. Viviana Risca.

ATACseq: GSE295258 (Reviewer token: ojihqayqnhethgj)

RNA-seq: GSE295260 (Reviewer token: qbsxqqouzxufzub)

CUT&Tag: GSE295261 (Reviewer token: shapgsuwfnqthcp)

## ACKNOWLEDGMENTS

L. Y. acknowledges support from a Natural Sciences and Engineering Research Council of Canada Postgraduate Scholarships – Doctoral scholarship. J. R. was supported by the American Cancer Society – Fairfield Research Council– Postdoctoral Fellowship, PF-23-1034949-01-CCB, https://doi.org/10.53354/ACS.PF-23-1034949-01-CCB.pc.gr.168157. L. A. W. acknowledges support from the NSF Graduate Research Fellowship Program (1946429). This work was supported in part by a V Scholar Award (V2019-011) from the V Foundation for Cancer Research, by a Starr Cancer Consortium Grant (I16-0058), a Scholar Award from The Rita Allen Foundation (S-2108-02147), and a Career Scientist Award from the Irma T. Hirschl/Monique Weill-Caulier Trust.

We acknowledge support from the Rockefeller University Genomics Resource Center (RRID:SCR_020986), and the High-Performance Computing Resource Center (RRID:SCR_025889). We thank Andrew Koff for providing us with the LS8817 cell line and for scientific insights. We thank Andrew Scortea for administrative support and his work in setting up the Risca laboratory. We thank Wei Zhou and Rochelle Shih for technical assistance. We thank all members of the Risca and Koff laboratories, Hironori Funabiki, Lydia Finley and Junyue Cao for insightful discussions and advice.

## AUTHOR CONTRIBUTIONS

J. L. Y.: Conceptualization of project, performed experiments, data analysis, visualization and curation, manuscript writing and editing. J. R.: Performed experiments, data analysis, manuscript writing and editing, and provided mentorship. N. P.: Bioinformatics analysis. L. A. W..: Performed experiments, image analysis and manuscript editing. A. O.: Performed experiments. M. P.: Performed experiments and manuscript editing. I. D.: Bioinformatics analysis. R. H.: Bioinformatics analysis. B. T.: Performed experiments and image analysis. V. I. R.: Conceptualization of project, resources, project supervision, mentorship, funding acquisition, manuscript writing and editing.

