## Supplemental Materials for "CDK4/6 inhibition induces a DNA damage-independent senescence-associated secretory phenotype driven by delayed activation of NF-κB"

### Table of Contents:

**Quantitative and statistical analysis**

**Materials and methods of supplemental figures**

**Supplemental Figures 1-6**

**Supplemental Tables 1-2**

**Supplemental Data 1-5**

**Supplemental Fig. 1:** Global transcriptome and epigenome changes over time across drug treatments, related to Fig. 2

**Supplemental Fig. 2:** Predicted upstream regulators of drug-specific transcriptional responses, related to Fig. 2

**Supplemental Fig. 3:** Additional evidence of NF-κB activity in LS8817 and MCF7 cells, related to Fig. 3

**Supplemental Fig. 4:** Additional ATAC-seq and epigenomic data, related to Fig. 4

**Supplemental Fig. 5:** BAY-11-7082 treatment suppresses the upregulation of NF-κB target genes in LS8817 cells, related to Fig. 5

**Supplemental Fig. 6:** Upregulation of cell surface receptors upstream of NF- κB activation is found in CDK4/6i treatment, related to Fig. 6

**Supplemental Table 1** (Excel file)**:** Quality control metrics of ATAC-seq and RNA-seq, CUT&Tag/CUT&RUN samples.

**Supplemental Table 2** (Excel file)**:** List of primer sequences used to RT-qPCR.

**Supplemental Data 1** (Excel file)**:** Log2FC, *Padj* values and assigned k-means cluster of expressed genes. Genes annotated as SASP genes, cell cycle genes, p53 targets, NF-κB targets and E2F targets are also shown.

**Supplemental Data 2** (Excel file)**:** All GO enrichment and IPA predicted upstream regulators results for RNA Clusters

**Supplemental Data 3** (Excel file)**:** IPA predicted activation scores of upstream regulators

**Supplemental Data 4** (Excel file)**:** Motif enrichment results on dynamic ATAC-seq peaks relative to Cycling or day 3 controls within drug treatment

**Supplemental Data 5** (Excel file)**:** Motif enrichment results of nearby dynamic ATAC-seq peaks to RNA Clusters of significantly changing genes

### QUANTIFICATION AND STATISTICAL ANALYSIS

#### Sample distance and PCA for sequencing data

Distance measures between samples were calculated by using the `dist` function on the transposed VST-normalized counts matrix for ATAC-seq or rlog-normalized counts matrix for RNA-seq in R. Prior to this, technical replicates were merged into biological replicates for ATAC-seq. The resulting distance values were plotted using pheatmap. Principal component analysis (PCA) plots were generated using DESeq2's `plotPCA` function.

#### RNA-sequencing pre-processing

Reads containing adapters, N > 10% (N represents the base cannot be determined) and low quality (Qscore<= 5) reads were filtered out. Pseudoalignment and quantification of transcripts were performed with kallisto (Bray et al. 2016).

#### Differential gene expression analysis

*LS8817:* Downstream analysis was performed in RStudio v3.6.3. Kallisto counts were imported into R using tximport v1.14.2, and differential gene expression analysis was conducted with DESeq2 1.26.0 (Love et al. 2014). Genes with fewer than 50 total counts across all samples were filtered out. Significantly differentially expressed genes were identified using a likelihood ratio test ($Padj < 0.05$) and a Wald test (absolute log2FC ≥ 2) relative to Cycling or day 3 treatment controls. A total of 3,744 differentially expressed genes were identified and clustered into seven RNA expression groups using k-means clustering. To characterize expression patterns across treatments, genes were further categorized as "Shared-down," "Palbo-upregulated," "Doxo-upregulated," or

"Shared-up." rlog-normalized gene expression values were visualized using the pheatmap R package, with values scaled by row to enhance comparability.

*MCF7:* Quantification of transcripts and read alignment were performed with STAR 2.7.10a Genes with at least 1 sample with > 5 counts were kept. Differential gene expression analysis was conducted with DESeq2 (Love et al. 2014) and significantly differentially expressed genes were identified using a likelihood ratio test (Padj < 0.05)

**Identification of SASP and transcription factor target genes**

***SASP genes:*** We used manually curated databases (Wang et al. 2022; Gleason et al. 2024) and the SASPAtlas (Basisty et al. 2020) (511 genes). Filtering for significantly changing genes present in all three databases, we identified 277 SASP genes that were associated with either therapy-induced senescence.

***Cell cycle genes:*** Significantly changing genes were annotated as Cell Cycle if they were found in one of the following databases (892 genes). Hallmark: “E2F TARGETS,” “G2M_CHECKPOINT;” Reactome: “CELL_CYCLE,” “CELL_CYCLE_CHECKPOINTS,” “CELL_CYCLE_MITOTIC;” KEGG: “CELL_CYCLE.”

***NF-κB target genes***: We used the Gilmore Lab NF-κB target gene database (https://www.bu.edu/nf-kb/gene-resources/target-genes/), the "TNFA SIGNALING VIA NFKB" Hallmark term, and Ingenuity Pathway Analysis (IPA) upstream regulators containing the pattern, "NFKB" (687 genes). Filtering for significantly changing genes present in all three sources, we identified 163 upregulated NF-κB target genes across RNA clusters.

### GO enrichment analysis

GO enrichment was performed using enricher function from clusterProfiler package (Yu et al. 2012), using gene IDs from each RNA cluster examining MSigDB databases (Liberzon et al. 2011; Liberzon et al. 2015). Default settings of *P*-value cutoff = 0.05 and *P*-adjust method = “BH” were used. Background gene set for GO enrichment was set as all genes expressed.

### GSEA analysis

Genes were ranked based on the product of log2FC*-log10(*P*-value), from DESeq2 results. GSEA was performed on the ranked list of genes using the GSEA function from clusterProfiler package v3.14.3 using the default settings.(Subramanian et al. 2005) Running enrichment score plots were generated using the gseaplot function with the setting “by = runningScore.”

### Ingenuity Pathway Analysis

To determine predicted upstream regulators, data were analyzed with the use of QIAGEN IPA (QIAGEN Inc., https://digitalinsights.qiagen.com/IPA) where the core analysis was performed on gene IDs from each RNA cluster (Krämer et al. 2014).

For all analyses, filtered upstream regulators were cross-referenced with a list of expressed genes in this cell line to ensure biological relevance. -log10(*Padj*) values for the top 5-10 enriched upstream regulators or activation scores for each condition were plotted as a heatmap using pheatmap R package.

**ATAC-sequencing preprocessing**

Adapters were trimmed using cutadapt and reads were aligned with bowtie2 to the hg38 genome. Following this, duplicate reads and reads mapping to the mitochondrial genome were removed. BAM files were converted to bedpe files, which were then converted to tagAlign files for MACS2 to call peaks on, areas of the genome enriched for aligned sequencing reads, indicating hyper-accessibility. MACS2 parameters were `-g 2.7e9 -p 0.2 --shift -75 --extsize 150 --nomodel -B --SPMR --keep-dup all --call-summits`. Reproducible peaks across technical replicates per condition were kept using Irreproducible Discovery Rate.(Li et al. 2011) A master nonredundant peak set was generated using a custom iterative peak filtering script which selects the peak with the lowest q-value generated by MACS2 from a set of overlapping peaks. Peaks were fixed to 500 bp in length, with 250 bp extending from the summit identified by MACS2.(Zhang et al. 2008) In total we identified 215,671 non-redundant peaks. Fragment counts under peaks were counted using chromVAR's `getCounts` function (Schep et al. 2017).

Normalized bigwigs were generated from merged BAM files of biological replicates for each condition using the `bamCoverage` function from deepTools with the following command: `bamCoverage -b $INPUT -o $OUTPUT --binSize 5 --scaleFactor $SCALE --extendReads` where $INPUT is the merged BAM file and $OUTPUT is the resulting BigWig file. $SCALE is the same scale factor used to generate bigWigs for fragment ends.

**Differential ATAC-seq peak analysis**

Differential peak analysis was performed using DESeq2 with the Wald test, based on pairwise comparisons of each timepoint relative to Cycling or day 3 treatment controls. Among the 215,671 unique peaks in our non-redundant master peak set, we identified 49,107 upregulated and 42,164 downregulated peaks across both drug treatments. Peaks were considered significant based on a Bonferroni-corrected *Padj* threshold (0.01/215,671 total peaks) and an absolute log2FC > 1 at any timepoint compared to Cycling or day 3 treatment controls. Upregulated peaks were categorized as "Doxo-upregulated" if they were significant in Doxo but not Palbo, "Palbo-upregulated" if significant in Palbo but not Doxo, and "Shared-upregulated" if significant in both treatments. The same criteria were applied for categorizing downregulated peaks.

To visualize read counts under significantly upregulated peaks, reads were variance-stabilizing transformed (VST), scaled by row and ordered based on their classification as Palbo-upregulated, Shared-upregulated, or Doxo-upregulated. Hierarchical clustering was applied within each drug category to improve the coherence of the heatmap structure. The same was performed for downregulated peaks.

**ATAC-seq Peak Annotation**

Significantly changing peaks at each timepoint relative to Cycling or day 3 treatment controls within each drug were merged into a single peak set. Peaks were annotated using the `annotatePeak` function from ChIPseeker with the UCSC hg38 knownGene TxDb (Yu et al. 2015).

**CUT&Tag and CUT&RUN pre-processing**

*CUT&Tag:* Sequencing data were processed using a modular pipeline implemented in Nextflow (https://github.com/riscalab/NEXDEP-Nextflow_DNA_Epigenomic_Pipeline). Quality control was assessed using FastQC, and reports were aggregated using MultiQC to generate summary metrics across all samples. Reads were aligned to the reference genome using **BWA (Burrows–Wheeler Aligner).** SAM files generated from alignment were converted to sorted and indexed BAM files using SAMtools. Alignment statistics, including mapping rates and read counts, were calculated for each sample. Reads overlapping ENCODE blacklist regions were removed. Filtered BAM files were re-indexed prior to downstream analysis.

*CUT&RUN:* Cutadapt was used to trim adapters using the following code: `cutadapt -a "CTGTCTCTTATACACATCTCCGAGCCCACGAGAC" -a "CTGTCTCTTATACACATCTGACGCTGCCGACGA" -o "$outfile1" -p "$outfile2" "$infile1" "$infile2"`. Reads with aligned to hg38 using bowtie2 using the following parameters `--end-to-end --very-sensitive --no-mixed --no-discordant --phred33 -I 10 -X 700`.

Peaks were called with SEACR software (Meers et al. 2019) with parameters specified to non-normalized and stringent threshold at 0.1 for H3K27ac and 0.01 for macroH2A1. Using bedtools, bam files were converted to bedpe files which were then converted to bedgraph format. Bedgraph files were used as input into SEACR peak caller. `bedtools intersect` was used to keep peaks that were reproducible in at least 2 out of 3 technical replicates before filtering for reproducible peaks across biological replicates. `Bedtools merge` was then used to generate a non-redundant master peak set across all conditions for each histone modification. FeatureCounts was used to quantify read pairs overlapping peaks from master peak sets across conditions, with the “count all overlapping features” option enabled, using the same code for CUT&Tag.

*Differential H3K27ac peak analysis*: Peaks that had at least 6 samples with normalized read counts >=5 were kept. DESeq was performed on 112,034 H3K27ac peaks. The significance criteria for all H3K27ac peaks were a *Padj* value of $< 0.05$ and an absolute log2FC value >=1 for the following conditions: Palbo vs. Cycling, day 14 Doxo vs. Cycling, Palbo vs. day 14 Doxo.

*Differential macroH2A1 peak analysis:* Peaks with fewer than 10 normalized read counts across all samples were filtered out. DESeq was performed on 184,456 macroH2A1 peaks. Significance criteria for all mH2A1 peaks was absolute log2FC value >=1 and *Padj* $< 0.01$ for the following comparisons: day 28 Palbo vs Cycling, day 14 Doxo vs Cycling, day 28 Palbo vs day 14 Doxo.

**Drug-activated enhancers identification**

Drug-activated enhancers were defined as regions with increased chromatin accessibility (ATAC-seq) overlapping regions of increased H3K27ac signal. All upregulated ATAC-seq peaks, regardless of treatment condition, were intersected with H3K27ac peaks increased in palbociclib, doxorubicin, or both treatments and classified as Palbo-, Doxo-, or Shared-upregulated enhancers, respectively.

**Nearby enhancers and macroH2A1-containing regions identification**

Genomic coordinates of genes were obtained from the comprehensive gene annotation file in GENCODE Release 41 (GRCh38.p13) (Frankish et al. 2019). For each RNA cluster of significantly changing genes, gene boundaries were extended by 50 kb upstream and downstream. Regions were then filtered to retain those overlapping with the extended genomic coordinates of genes identified in shared-upregulated gene module (Fig. 2D). Additionally, nearby static peaks (peaks that did not meet the significance criteria) were identified using the same approach.

#### Motif enrichment analysis

Motif enrichment analysis was performed using Analysis of Motif Enrichment (AME) from the MEME SUITE v5.0.2 (McLeay and Bailey 2010). FASTA sequences under regions of interest were extracted using the Biostrings R package (Pages et al. 2013). Enriched motifs were identified using the HOCOMOCO v11 database (Kulakovskiy et al. 2018) with the following command: `ame --verbose 1 --scoring avg --method fisher --hit-lo-fraction 0.25 --evalue-report-threshold 10.0 --control $CONTROL --kmer 2 $INPUT HOCOMOCOv11_core_HUMAN_mono_meme_format.meme`.

Here `$CONTROL` represents the FASTA file containing sequences from background peaks, and `$INPUT` contains sequences from foreground peaks.

*MEME-AME on nearby drug activated enhancers:* Drug activated enhancers were first defined as regions showing increased accessibility (ATAC-seq) and H3K27ac signal upon treatment. Among these, enhancers located within ±50 kb of shared-upregulated genes (Fig. 2D), including those overlapping gene bodies, were selected. Static ATAC-seq peaks within the same ±50 kb regions served as the background set.

#### Coverage plots

Bams across all technical and biological replicates were merged and sorted by coordinate using samtools. Merged RPGC-normalized bigwigs for each condition was generated using deepTools `bamCoverage` function with a bin-size of 10 and effectiveGenomeSize of 2913022398. RPGC-normalized merged bigWig files for H3K27ac and were used as input to plot signal ± 1 kb of centers of ATAC-seq peaks that were considered as drug-activated enhancers. RPGC-normalized merged BigWig files for macroH2A1 was used as input to plot macroH2A1 signal 1 kb upstream and downstream of macroH2A1 peak centers. BigWig signal was plotted within 1 kb of motifs using deepTools with the `computeMatrix` function (`--referencePoint center`) and `plotHeatmap` function.

### MATERIALS AND METHODS OF SUPPLEMENTAL FIGURES

#### Drug treatment and RNA-seq on MCF7 cells

MCF7 cells were treated with 100 nM doxorubicin (treated for 48 hours, followed by drug washout), 1 μM palbociclib or 100 nM palbociclib + 10 nM fulvestrant. Cells were harvested at day 14, day 21 and day 18 respectively, since treatment started. RNA extraction was done using the Quick-RNA Microprep Kit (Cat# R1050, Zymo Research, Irvine, California). Total RNA was sent to PlasmidSaurus for single-end RNA sequencing.

*Preprocessing by Plasmidsaurus:* Quality of the fastq files was assessed using FastQC v0.12.1. Reads were then quality filtered using fastp v0.24.0 with poly-X tail trimming, 3' quality-based tail trimming, a minimum Phred quality score of 15, and a minimum length requirement of 50 bp PCR and optical duplicates were removed using UMI-based deduplication with UMIcollapse v1.1.0. Alignment quality metrics, strand specificity, and read distribution across genomic features were assessed using RSeQC v5.0.4 and Qualimap v2.3, with results aggregated into a comprehensive quality control report using MultiQC v1.32.

#### Identification of transcription factor target genes

***P53 target genes:*** Significantly changing genes were cross-referenced with the curated p53 target list from Fischer and the "P53 PATHWAY" Hallmark term (473 genes), identifying 102 upregulated p53 target genes across RNA clusters.

***E2F target genes***: We filtered significantly changing genes against the “E2F TARGETS” Hallmark term and Ingenuity Pathway Analysis (IPA) upstream regulators containing the pattern, “E2F” (291 genes), identifying 164 downregulated E2F target genes.

**Activation scores of predicted upstream regulators**

To determine the activation scores and visualize predicted relationships between upstream regulators, IPA core analysis was performed on log2FC values of gene IDs across timepoints relative to Cycling or day 3 treatment controls with a threshold set to an absolute value of log2FC = 1.

To identify differentially activated upstream regulators and compare between drugs, an IPA comparison was performed on the individual core analyses results, generating a matrix of activation scores. To filter for the top 10-12 most differentially activated upstream regulators, the sum of activation scores across timepoints within each drug was calculated. Upstream regulators with activation scores significantly higher in one treatment condition relative to the other were selected based on greatest magnitude differences between drugs. Conversely, upstream regulators significantly activated in Doxo-treated conditions but repressed in Palbo-treated conditions were selected based on an analogous threshold.

**Motif enrichment analysis**

*Upregulated ATAC-seq peaks:* All upregulated ATAC-seq peaks across timepoints were merged into a single peakset and were subsequently classified as Palbo-Up (palbociclib-specific), Doxo-Up (doxorubicin-specific), or Shared-Up (upregulated in both treatments).

For each group, motif enrichment was performed using the union of upregulated peaks from the other conditions as the background. Specifically, Palbo-Up peaks were compared against Shared-Up and Doxo-Up peaks combined, and the same strategy was applied for the other groups. FASTA sequences were extracted from upregulated peaks at each time point relative to either Cycling (untreated) or day 3-treated controls within each drug treatment; these were used as the foreground peak set, with the corresponding downregulated ATAC-seq peaks serving as the background set. The same approach while switching the foreground and background sets was used to identify motifs enriched in downregulated ATAC-seq peaks.

*Nearby drug-activated enhancers*: Up- or down-regulated nearby peaks for each drug treatment were used as the foreground set, while static nearby peaks served as the background set.

**Coverage plots**

*ATAC-seq accessibility signal plots around motifs*: Tn5 insertion sites were identified as fragment ends from merged BAM files of biological replicates for each condition and saved as bigWig files. This was performed in R using a custom script. In brief, the `resize` and `shift` functions in GenomicRanges were used to retain and shift the ends of reads to +4, -5 bp to account for Tn5 insertion bias. The `coverage` function was used to convert the fragment ends into a RleList object. To normalize bigWig files, reads were counted in 500 bp bins tiled across the genome, and the resulting count matrix was used in DESeq to calculate size factors with the `estimateSizeFactors` function. Normalization was performed by multiplying the coverage by a scale factor (1/DESeq size factor) before exporting the resulting object as a bigWig.

Motif genomic coordinates were identified using position weight matrices from HOCOMOCO v11 and scanned across the hg38 genome with motifmatchr 1.8.0 R package (Kulakovskiy et al. 2018). Motifs overlapping up-regulated ATAC-seq peaks at any timepoint for each drug category (Doxo-upregulated, Shared-upregulated, Palbo-upregulated) were saved as BED files and used for coverage plotting in deepTools (Ramírez et al. 2016). Tn5 insertion site coverage was plotted within 1 kb of motifs using deepTools with the `computeMatrix` function (`--referencePoint center`) and `plotProfile` or `plotHeatmap` function.

### SUPPLEMENTAL FIGURES

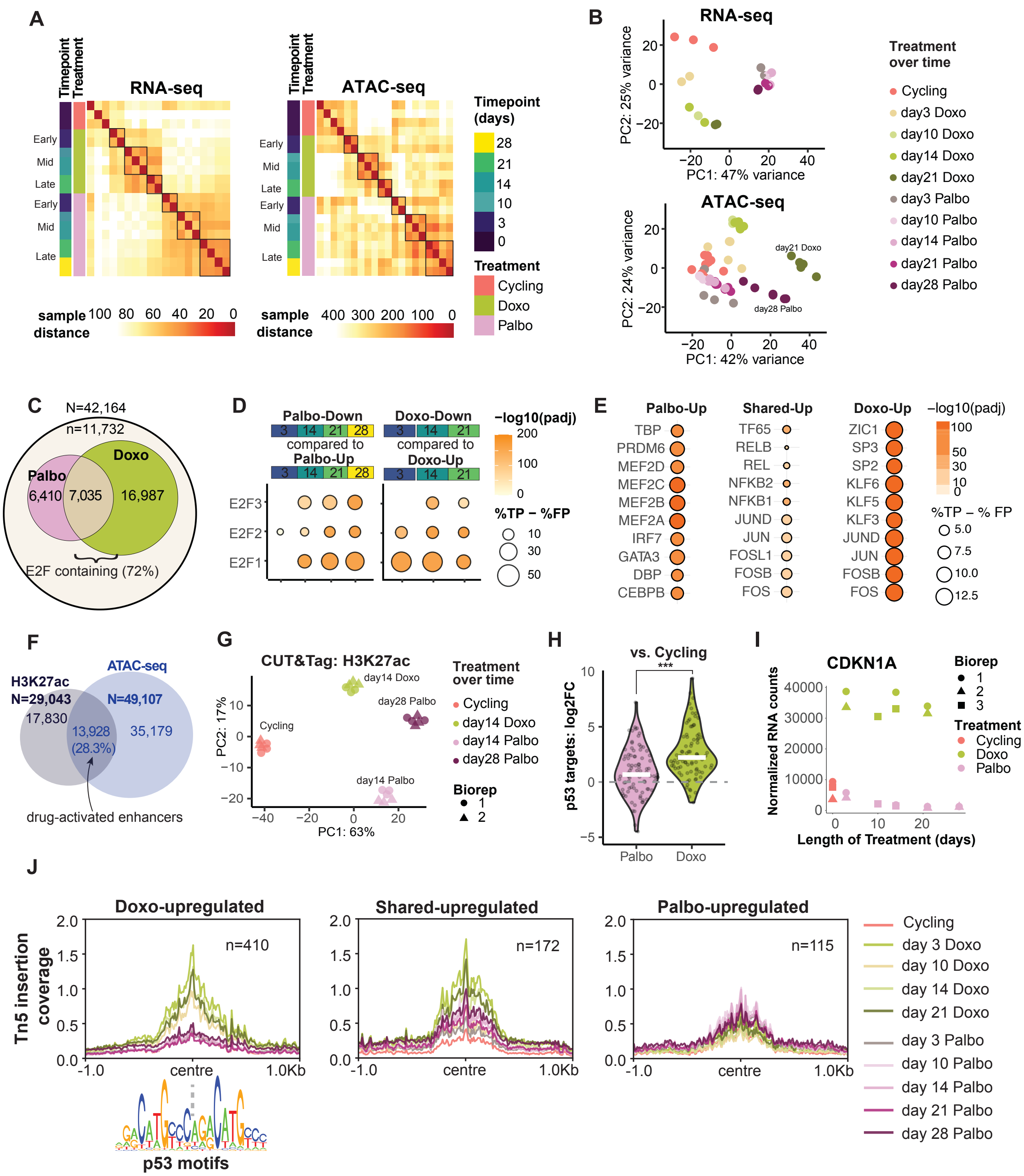

**Supplemental Fig.1: Global transcriptome and epigenome changes over time across drug treatments, related to Figure 2**

A. Distance measures between samples ordered by treatment and timepoint for RNA-seq (left) and ATAC-seq (right).
B. Principal component analysis on RNA-seq (top) and ATAC-seq (bottom) samples.
C. Venn diagram showing the number of peaks with decreased accessibility that contain E2F family of motifs relative to all downregulated ATAC-seq peaks
D. Motif enrichment results of E2F family of motifs in downregulated ATAC-seq peaks relative to upregulated peaks (based on differential accessibility relative to Cycling) within each drug. The timepoints indicated represent the peak sets chosen as input for MEME-AME. Dot intensity represents -log10(Padj) values, while dot size reflects the difference between true positive and false positive peaks (%TP - %FP).
E. Motif enrichment results of top 10 most enriched motifs in ATAC-seq peaks upregulated in palbociclib (Palbo-Up: left), both treatments (Shared-Up: middle) or doxorubicin (Doxo-Up: right), merged across all timepoints relative to Cycling and day 3, using unique upregulated peaks from the alternate condition as background.
F. Venn diagrams showing the number of upregulated H3K27ac peaks that overlap with upregulated ATAC-seq peaks, defined as drug-activated enhancers (see Methods). Color of text indicates the number of peaks from each peak set (H3K27ac: blue-grey; ATAC-seq: dark blue).
G. Principal component analysis of H3K27ac CUT&Tag samples.
H. Log2 fold change (log2FC) values across all significantly changing p53 target genes in day 14 Palbo and day 14 Doxo, relative to Cycling. Paired student t-test shows difference between Palbo and Doxo is significant, with P-value = 4.468x10-8.
I. Normalized RNA-seq counts of the p53 target gene, CDKN1A (p21) over time.
J. Tn5 insertion sites around aggregated p53 motifs overlapping Doxo-, Shared- or Palbo-upregulated ATAC-seq peaks relative to Cycling (peaks from Figure 2A). The number of overlapping peaks with p53 motifs per condition is shown.

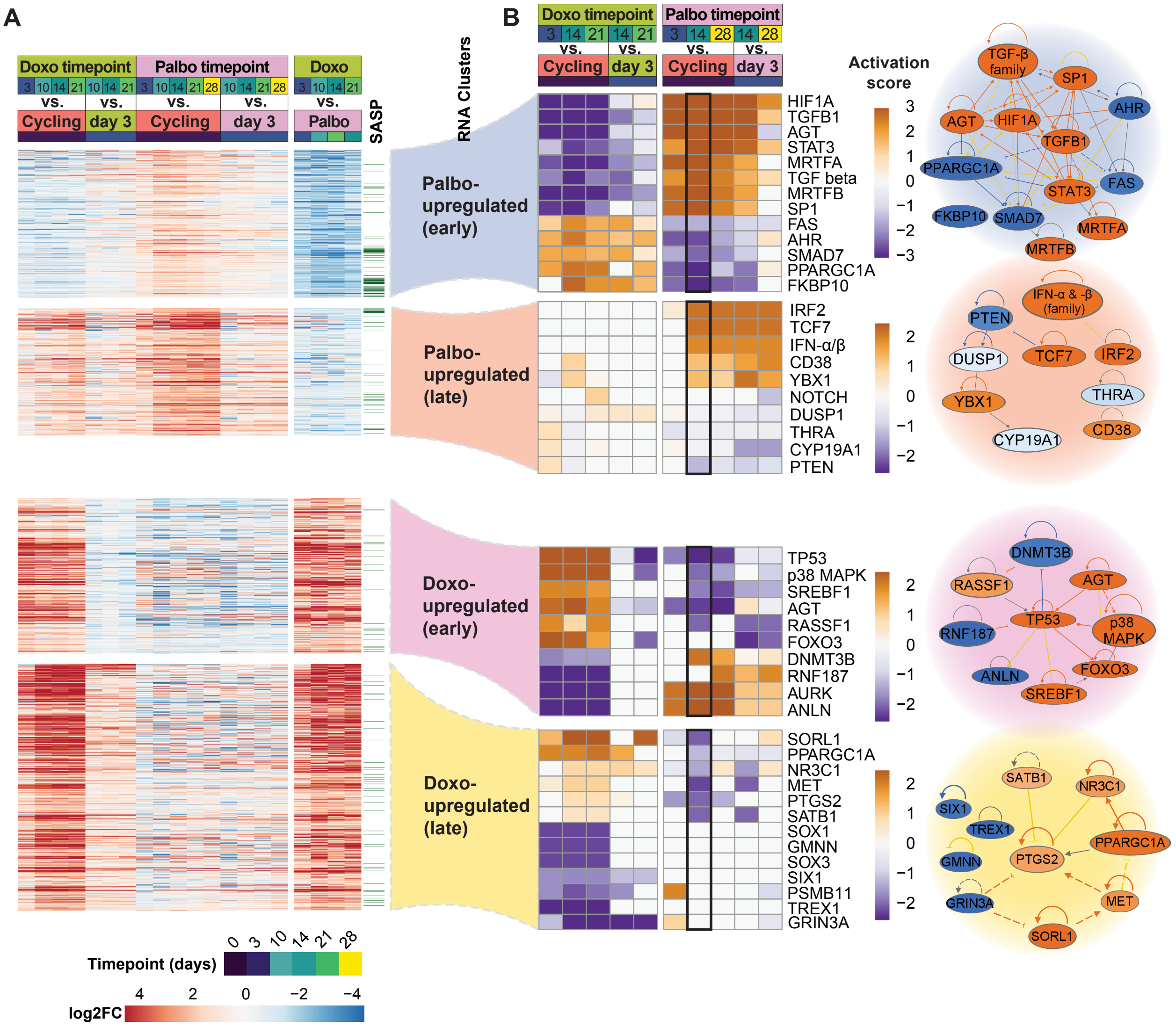


### Supplemental Fig.2: Predicted upstream regulators of drug-specific transcriptional responses, related to Figure 2

A. Log2 fold change (log2FC) values of differentially upregulated genes from Figure 2D, across different timepoints (days of treatment) and drug conditions, relative to Cycling or day 3 controls for each drug treatment.

B. Activation scores of the most differentially activated upstream regulators between Doxo and Palbo, corresponding to drug-distinct clusters of genes from Figure 2D. Differentially activated regulators were identified by summing activation scores across timepoints within each treatment and ranking them from highest to lowest (see Methods). A black outline marks the timepoint used to generate networks from IPA core analysis. Predicted relationships are shown (right) for each RNA cluster with legend at the bottom-right indicating the confidence and type of predicted relationships.

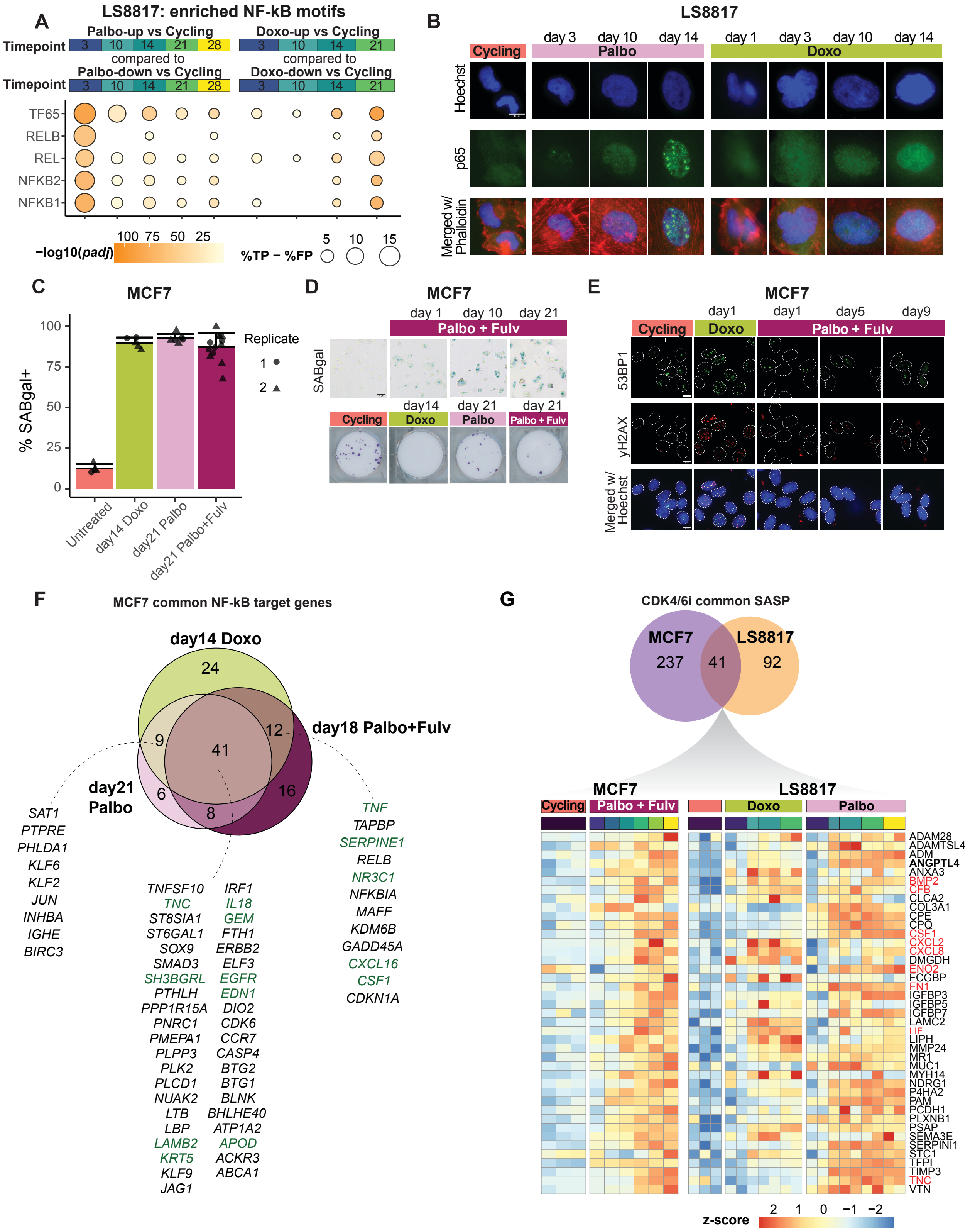
A
LS8817: enriched NF-kB motifs
Timepoint
Palbo-up vs Cycling
3 10 14 21 28
compared to
Palbo-down vs Cycling
Doxo-up vs Cycling
3 10 14 21
Doxo-down vs Cycling
TF65
RELB
REL
NFKB2
NFKB1
−log10(padj)
100 75 50 25
%TP − %FP
5 10 15
B
LS8817
Cycling
Palbo
Doxo
day 3 day 10 day 14
day 1 day 3 day 10 day 14
Hoechst
p65
Merged w/ Phalloidin
C
MCF7
% SABgal+
100 75 50 25 0
Replicate
1
2
Untreated
day14 Doxo
day21 Palbo
day21 Palbo+Fulv
D
MCF7
day 1 day 10 day 21
Palbo + Fulv
SABgal
day14 day 21 day 21
Cycling
Doxo
Palbo
Palbo + Fulv
E
MCF7
day1 day1 day5 day9
Cycling
Doxo
Palbo + Fulv
53BP1
yH2AX
Merged w/ Hoechst
F
MCF7 common NF-kB target genes
day14 Doxo
day18 Palbo+Fulv
day21 Palbo
24 12 9 41 6 16 8
SAT1 PTPRE PHLDA1 KLF6 KLF2 JUN INHBA IGHE BIRC3
TNFSF10 TNC ST8SIA1 ST6GAL1 SOX9 SMAD3 SH3BGRL PTHLH PPP1R15A PNRC1 PMEPA1 PLPP3 PLK2 PLCD1 NUAK2 LTB LBP LAMB2 KRT5 KLF9 JAG1
IRF1 IL18 GEM FTH1 ERBB2 ELF3 EGFR EDN1 DIO2 CDK6 CCR7 CASP4 BTG2 BTG1 BLNK BHLHE40 ATP1A2 APOD ACKR3 ABCA1
TNF TAPBP SERPINE1 RELB NR3C1 NFKBIA MAFF KDM6B GADD45A CXCL16 CSF1 CDKN1A
G
CDK4/6i common SASP
MCF7
LS8817
237 41 92
MCF7
LS8817
Cycling
Palbo + Fulv
Doxo
Palbo
ADAM28 ADAMTSL4 ADM ANGPTL4 ANXA3 BMP2 CFB CLCA2 COL3A1 CPE CPQ CSF1 CXCL2 CXCL8 DMGDH ENO2 FCGBP FN1 IGFBP3 IGFBP5 IGFBP7 LAMC2 LIF LIPH MMP24 MR1 MUC1 MYH14 NDRG1 P4HA2 PAM PCDH1 PLXNB1 PSAP SEMA3E SERPINI1 STC1 TFPI TIMP3 TNC VTN
z-score
2 1 0 −1 −2

**Supplemental Fig.3: Additional evidence of NF-κB activity in LS8817 and MCF7 cells, related to Figure 3**

A. Motif enrichment results of NF-κB family of motifs in up-regulated ATAC-seq peaks relative to down-regulated peaks (based on differential accessibility relative to Cycling) within each drug. The timepoints indicated represent the peak sets chosen as input for MEME-AME. Dot intensity represents -log10(*p-adj*) values, while dot size reflects the difference between true positive and false positive peaks (%TP - %FP).
B. Representative images of p65 immunofluorescence signal in LS8817 cells taken at 60x objective magnification throughout therapy-induced senescence. Cells were treated with 1uM palbociclib (Palbo) continuously or 48 hours with 100nM doxorubicin (Doxo) before wash out until harvest. The scale bar is 15 μm.
C. Percent nuclei counted that had cellular SA-βgal+ staining in MCF7 cells for the following conditions: doxorubicin (Doxo), 1 μM palbociclib (Palbo), 100 nM palbociclib + 10 nM fulvestrant (Palbo + Fulv). Number of nuclei counted was > 250 per condition at 20x objective magnification. Each point represents a field of view, with different shapes indicating biological replicates. Error bars denote standard deviation.
D. Crystal violet staining after 17 days of clonogenic outgrowth from 400 MCF7 cells seeded.
E. Representative images showing different levels of DNA damage (γH2AX foci & 53BP1 foci) in Doxo and Palbo+Fulv treated MCF7 cells over time. The scale bar is 10 μm.
F. Venn diagram showing overlap of upregulated NF-κB target genes across conditions relative to untreated controls in MCF7. Gene IDs are indicated, with SASP genes highlighted in dark green.
G. Top: Venn diagram showing overlap of upregulated CDK4/6-induced SASP genes common across LS8817 and MCF7 cells. Z-scores from r-log normalized counts of CDK4/6-induced SASP genes commons across cancer cell types. SASP genes in red are also annotated as NF-κB target genes.

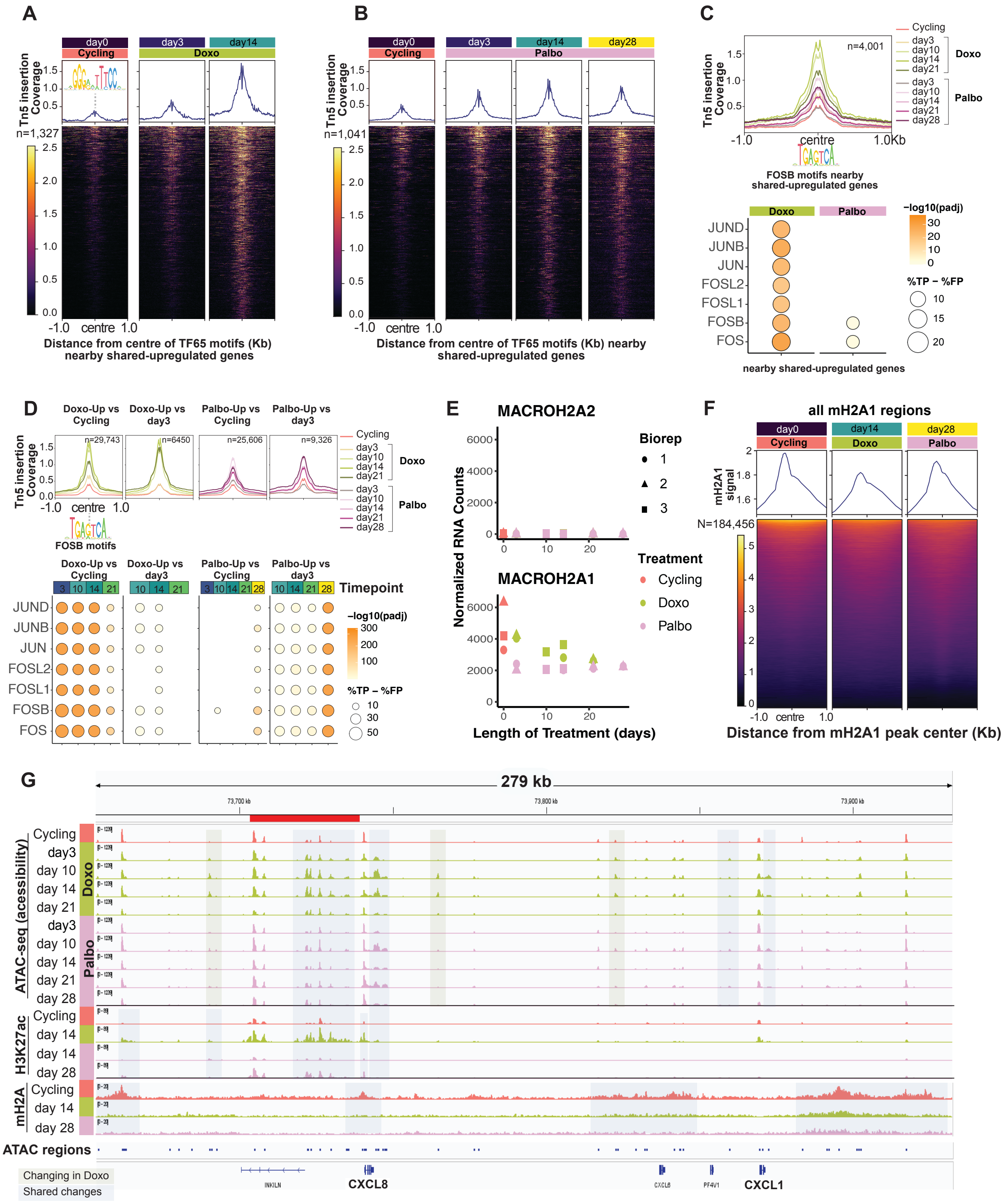
A
B
C
D
E
F
G
Distance from centre of TF65 motifs (Kb) nearby shared-upregulated genes
FOSB motifs nearby shared-upregulated genes
nearby shared-upregulated genes
FOSB motifs
Timepoint
MACROH2A2
MACROH2A1
Normalized RNA Counts
Length of Treatment (days)
Biorep
Treatment
Cycling
Doxo
Palbo
all mH2A1 regions
Distance from mH2A1 peak center (Kb)
279 kb
ATAC-seq (acessibility)
H3K27ac
mH2A
ATAC regions
CXCL8
CXCL1
Changing in Doxo
Shared changes

**Supplemental Fig.4: Additional ATAC-seq and epigenomic data, related to Figure 4**

A. Tn5 insertion site coverage at the centre of NF-κB subunit, TF65 (p65) motifs overlapping nearby Doxo-upregulated ATAC-seq peaks, defined as within 50kb of shared-upregulated genes. Number of nearby peaks is shown on the left. Scale bar represents bigwig signal of normalized coverage.
B. Same as A, but for nearby Palbo-upregulated ATAC-seq peaks.
C. **Top:** Tn5 insertion site coverage at the centre of the AP-1 family member, FOSB motifs overlapping all upregulated ATAC-seq peaks, in either drug treatment, within 50kb of shared-upregulated genes. **Bottom:** Motif enrichment results of AP-1 family of motifs (JUN/FOS) in upregulated ATAC-seq peaks within 50kb of shared-upregulated genes (based on differential accessibility relative to Cycling or day 3) relative to nearby static peaks for each drug. The timepoints indicated represent the peak sets chosen as input for MEME-AME. Dot intensity represents -log10(Padj) values, while dot size reflects the difference between true positive and false positive peaks (%TP - %FP).
D. **Top:** Tn5 insertion site profiles around FOSB motifs overlapping all ATAC-seq peaks. ATAC-seq peaks increasing in accessibility at any timepoint in response to Doxo or Palbo, compared to Cycling or day 3 controls within each drug. The number of overlapping peaks with FOS motifs is shown. **Bottom:** Motif enrichment results of AP-1 family of motifs in upregulated ATAC-seq peaks at each timepoint relative to Cycling or day 3 controls within each drug.
E. Normalized RNA-seq counts of *macroH2A2* and *macroH2A1* over time.
F. macroH2A1 (mH2A1) signal for all mH2A1 peaks called. The total number of mH2A1 peaks is shown on the left. Scale bar represents bigwig signal of normalized coverage.
G. Genome browser view of *CXCL1* and *CXCL8* loci. Normalized bigwig tracks for ATAC-seq, H3K27ac and macroH2A1 (mH2A) are shown. Upregulated regions occurring in both treatments are highlighted in blue (Shared changes) while Doxo-upregulated regions are highlighted in green (Changes in Doxo).

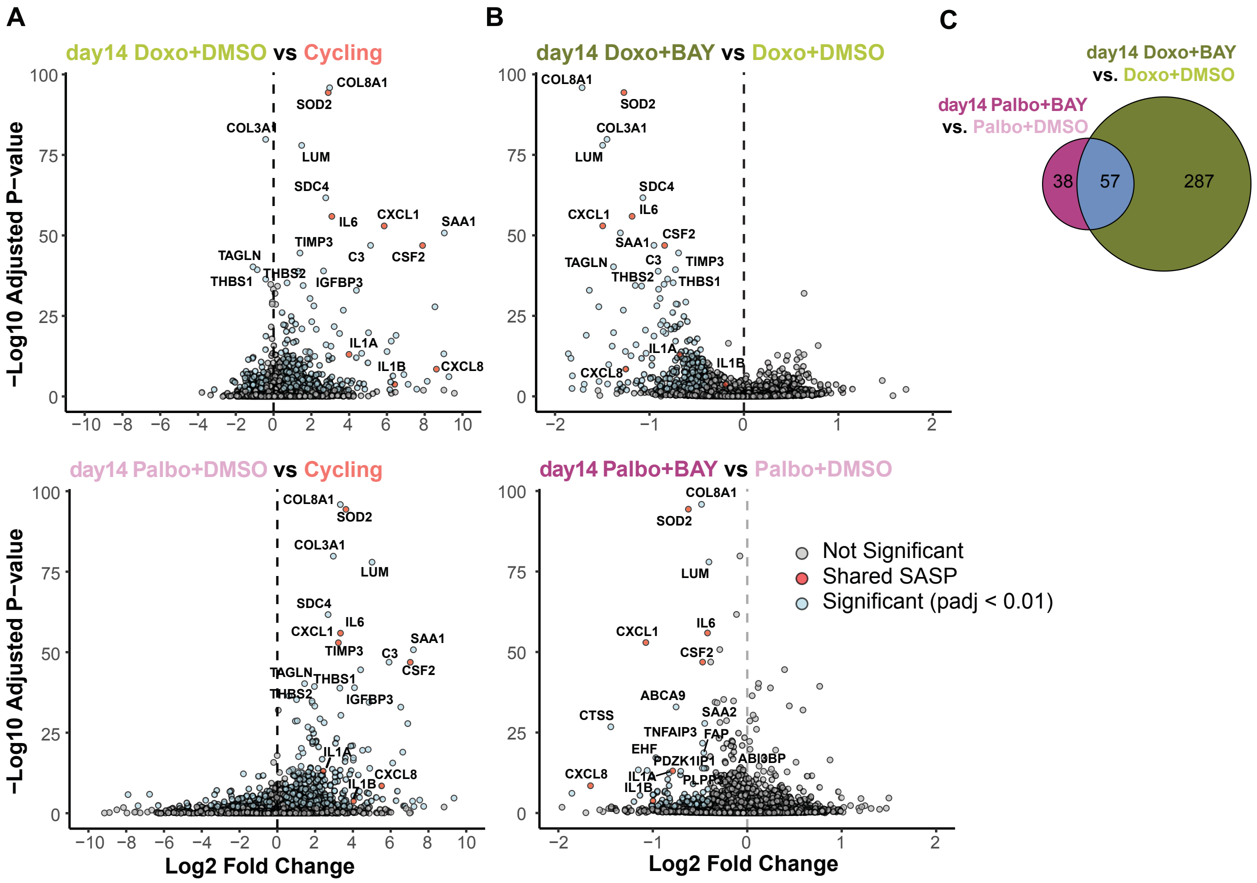


**Supplemental Fig.5: BAY-11-7082 treatment suppresses the upregulation of NF-κB target genes in LS8817 cells, related to Figure 5**

A. Volcano plot from RNA-seq analysis showing shared NF-κB-driven SASP genes and other top 15 most significantly changing genes in day 14 palbociclib or doxorubicin treated with DMSO control.
B. Same as (A) but for day 14 palbociclib or doxorubicin treated with BAY.
C. Venn diagram showing the number of genes downregulated by BAY relative to day 14 palbociclib or doxorubicin treated DMSO controls.

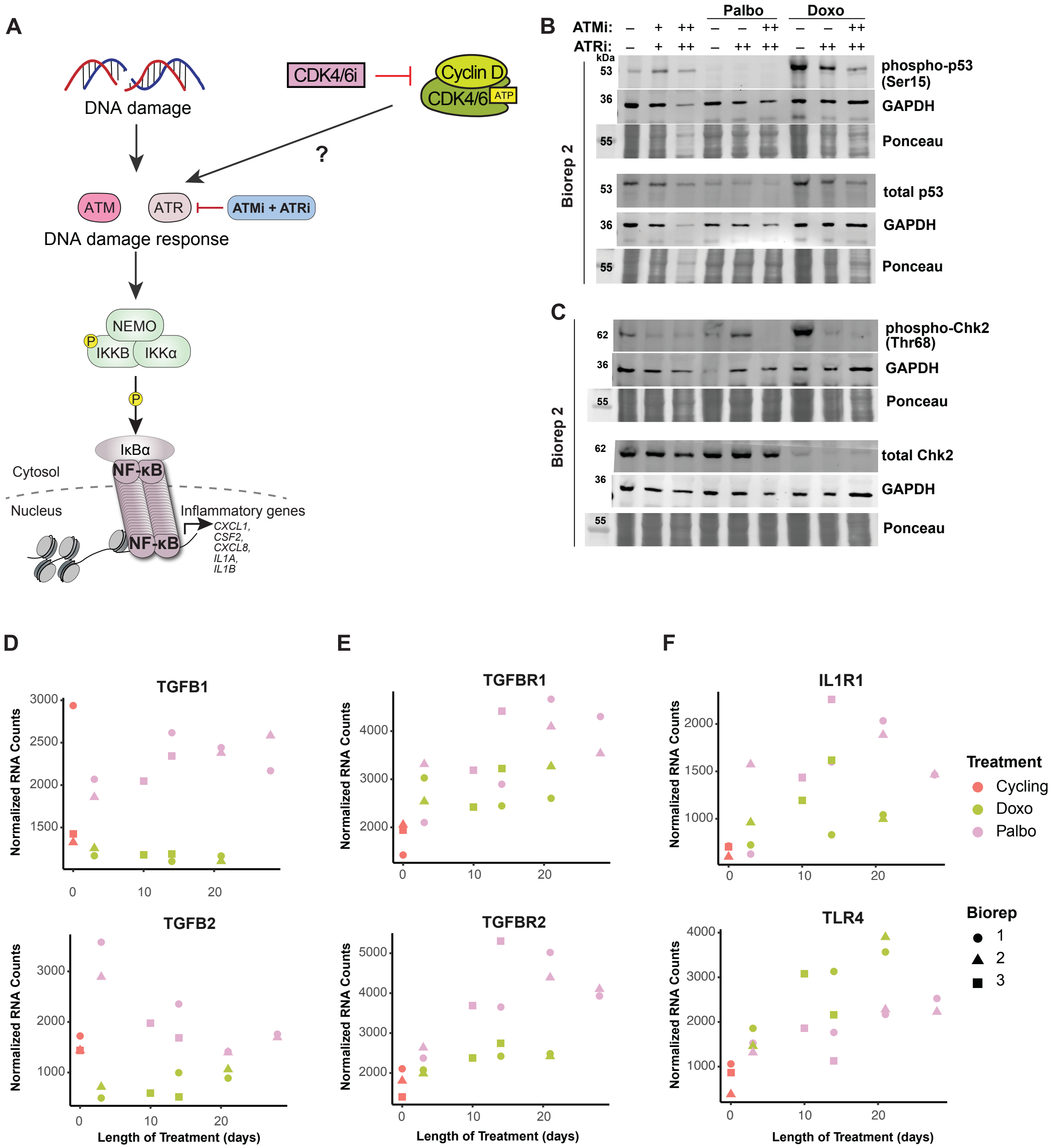

A
DNA damage
CDK4/6i
Cyclin D
CDK4/6
ATP
?
ATM
ATR
ATMi + ATRi
DNA damage response
NEMO
P
IKKB
IKKα
IκBα
NF-κB
Cytosol
Nucleus
Inflammatory genes
CXCL1, CSF2, CXCL8, IL1A, IL1B
B
Palbo
Doxo
ATMi:
ATRi:
kDa
Biorep 2
phospho-p53 (Ser15)
GAPDH
Ponceau
total p53
C
phospho-Chk2 (Thr68)
total Chk2
D
TGFB1
TGFB2
E
TGFBR1
TGFBR2
F
IL1R1
TLR4
Normalized RNA Counts
Length of Treatment (days)
Treatment
Cycling
Doxo
Palbo
Biorep
1
2
3

**Supplemental Fig.6: Upregulation of cell surface receptors upstream of NF- κB activation is found in CDK4/6i treatment, related to Figure 6**

A. Schematic of the experimental strategy testing whether inhibition of the DNA damage response kinases ATM and ATR affects NF-κB activation in CDK4/6i treatment in LS8817 cells.
B. Second biological replicate of Western blots for phospho-p53 and total p53, corresponding to Figure 6A.
C. As in (B), but for phospho-Chk2 and total Chk2.
D. Normalized RNA-seq counts of *TGFB1* and *TGFB2* over time.
E. Normalized RNA-seq counts of *TGFBR1* and *TGFBR2* over time.
F. Normalized RNA-seq counts of *IL1R1* and *TLR4* over time.
